# Imbalance between RNA-binding proteins and the WAVE regulatory complex underlies the toxic gain-of-function effects of the *CYFIP2* Arg87Cys variant in neurodevelopmental disorders

**DOI:** 10.64898/2026.10.27.762134

**Authors:** U Suk Kim, Yukyung Jun, Hyewon Kim, Yeji Yang, Muwon Kang, Ruiying Ma, Hyae Rim Kang, Sungjin Ju, Hoibin Jeong, Su Yeon Kim, Yoonhee Kim, Jun Young Oh, Eunjoon Kim, Kyoungmi Kim, Hyojin Kang, Jin Young Kim, Kihoon Han

## Abstract

Pathogenic variants in cytoplasmic FMR1-interacting protein 2 (*CYFIP2*) cause neurodevelopmental disorders, with hotspot substitutions at arginine 87 (Arg87) associated with particularly severe symptoms. CYFIP2 functions both as a component of the WAVE (Wiskott–Aldrich syndrome protein family verprolin-homologous protein) regulatory complex (WRC), regulating the actin cytoskeleton, and as an interactor of RNA-binding proteins (RBPs) involved in mRNA processing/translation. Although Arg87 variants reduce CYFIP2 protein stability, heterozygous knock-in mice carrying Arg87Cys (*Cyfip2^+/R87C^*) show markedly more severe phenotypes than heterozygous knockout (*Cyfip2^+/-^*) mice, supporting a toxic gain-of-function mechanism. However, how the dual roles of CYFIP2 contribute to this mechanism remains unclear. To address this, we examined *Cyfip2^+/R87C^* mice at postnatal week 7, before overt neuronal loss and gliosis, focusing on the progressive dispersion of hippocampal cornu ammonis 1 (CA1) pyramidal neurons, a phenotype absent in *Cyfip2^+/-^*mice. Single-nucleus RNA sequencing suggested candidate alterations related to translation and cell adhesion/migration in CA1 pyramidal neurons. Complementarily, human embryonic kidney 293T cells expressing CYFIP2 Arg87Cys showed defects in proliferation and morphology, accompanied by altered expression of cell adhesion-related genes. Quantitative proteomic analysis of the CA1 region uncovered an imbalance between CYFIP2-associated protein networks, characterized by increased RBPs and reduced WRC components. Phosphoproteomic analysis further revealed altered phosphorylation of synaptic and translational regulators. Consistently, whole-cell recordings and biocytin-based morphological analyses showed increased frequency of miniature excitatory postsynaptic currents and altered dendritic morphology in *Cyfip2^+/R87C^* CA1 pyramidal neurons, most prominently in the more severely dispersed deep-layer neurons. Moreover, the cell adhesion molecule Eph receptor A3 was reduced, possibly as a result of RBP dysregulation. Together, these findings suggest that an imbalance between RBP- and WRC-associated pathways underlies the gain-of-function effects of CYFIP2 Arg87Cys and contributes to early neuronal phenotypes, most notably CA1 layer dispersion. This mechanism, distinct from *CYFIP2* haploinsufficiency, supports allele-specific therapeutic strategies for Arg87 variants.

## Introduction

Cytoplasmic FMR1-interacting protein 2 (CYFIP2) is an evolutionarily conserved protein that is highly expressed in the brain (Schenck et al., 2001). As a critical component of the Wiskott–Aldrich syndrome protein family verprolin-homologous protein (WAVE) regulatory complex (WRC), the primary function of CYFIP2 has been considered to be the regulation of neuronal actin polymerization and branching, processes that are essential for proper axonal and dendritic spine development (Chen et al., 2010; Cioni et al., 2018; Han & Ko, 2023; Y. Lee et al., 2017; Rottner et al., 2021). Intriguingly, however, recent proteomic studies have revealed that CYFIP2 can also physically and functionally interact with various RNA-binding proteins (RBPs) in the brain, suggesting additional roles in mRNA processing and translation. Specifically, the CYFIP2 interactome in the neonatal mouse forebrain comprises not only components of the WRC but also several RBPs, including Argonaute (Ago) and Pumilio (PUM) proteins (Y. Lee et al., 2020). Moreover, quantitative global proteomic analysis of embryonic *Cyfip2*-null (i.e., conventional knock-out [KO]) mouse forebrain revealed significant down-regulation of both WRC components and RBPs, particularly proteins associated with membraneless organelles (Zhang et al., 2024). Nevertheless, whether and how the balance between these dual functions of CYFIP2 is regulated under physiological and pathological conditions remains largely unknown (Ma, Kim, et al., 2025; Y. Zhang et al., 2019).

Since 2018, *de novo* variants of *CYFIP2* have been identified in pediatric patients with early-onset epileptic encephalopathy accompanied by developmental regression, classified as developmental and epileptic encephalopathies 65 (DEE65) (Begemann et al., 2021; Kim et al., 2025; Ma, Kim, et al., 2025; Nakashima et al., 2018; Zhong et al., 2019; Zweier et al., 2019). The clinical features include early-onset seizures, muscular hypotonia, intellectual disability, microcephaly, and gastrointestinal abnormalities. Notably, approximately 40% of reported cases harbor hotspot variants at the arginine 87 (Arg87) residue of CYFIP2, including Arg87Cys, Arg87Leu, and Arg87Pro. Patients with these variants, most of whom are diagnosed with West syndrome (also referred to as infantile spasms), exhibit more severe clinical manifestations than those carrying other *CYFIP2* variants (Begemann et al., 2021; Squire et al., 2025).

To better understand the molecular and cellular pathological mechanisms underlying the *CYFIP2* Arg87 hotspot variants, we previously generated and characterized a knock-in mouse line carrying the Arg87Cys variant in *Cyfip2* gene (*Cyfip2^+/R87C^* mice). Initial analysis of *Cyfip2^+/R87C^* mouse brains, combined with overexpression experiments in cultured cells, demonstrated that the Arg87Cys variant induces hyperubiquitination of the CYFIP2 protein, leading to its proteasomal degradation (Kang et al., 2023; Lee et al., 2019). Based on these findings, we initially reasoned that the Arg87 hotspot variants might function as loss-of-function alleles by reducing CYFIP2 protein levels, analogous to *CYFIP2* haploinsufficiency (Han et al., 2015; Kim et al., 2020; S. H. Lee et al., 2020).

To our surprise, however, a parallel long-term investigation on *Cyfip2^+/R87C^* mice and *Cyfip2^+/-^* mice (i.e., conventional heterozygous mice), a model of *CYFIP2* haploinsufficiency, revealed that *Cyfip2^+/R87C^* mice exhibited much more severe phenotypes than *Cyfip2^+/-^* mice (Ma, Kang, et al., 2025; Ma et al., 2026). Specifically, only *Cyfip2^+/R87C^* mice showed developmental regression, along with age-dependent spontaneous recurrent seizures and premature lethality. At the cellular level, synapse remodeling and neuronal loss, together with gliosis involving oligodendrocytes, microglia, and astrocytes, were observed in the hippocampus of adult *Cyfip2^+/R87C^* mice, but not in *Cyfip2^+/-^* mice. Moreover, bulk-tissue RNA sequencing from neonatal to adult stages revealed completely distinct trajectories of cortical and hippocampal gene expression between the two lines. Collectively, these results indicate that the Arg87Cys variant exerts toxic gain-of-function effects, rather than simply causing down-regulation of CYFIP2 proteins (Ma, Kang, et al., 2025; Ma, Kim, et al., 2025). Clinically, this distinction is critical, because a gain-of-function mechanism would call for allele-specific inhibitory strategies, such as antisense oligonucleotides (ASOs) targeting the Arg87Cys allele (Aartsma-Rus et al., 2026; Lauffer et al., 2024), whereas a loss-of-function mechanism would instead require approaches aimed at restoring CYFIP2 levels or function.

Despite this phenotypic evidence, the molecular mechanisms underlying the gain-of-function effects of Arg87Cys remain unknown. In our previous study, a longitudinal analysis of *Cyfip2^+/R87C^* mice revealed pronounced gliosis in the hippocampus, emerging around postnatal week 14 (PWK 14), which coincides with the onset of spontaneous recurrent seizures (Ma, Kang, et al., 2025). Although this represents a striking and specific cellular phenotype observed only in *Cyfip2^+/R87C^* mice and not in *Cyfip2^+/-^* mice, the fact that CYFIP2 protein is predominantly expressed in neurons rather than in glial cells suggests that this gliosis is likely a secondary consequence of earlier neuronal alterations (Ma et al., 2022; Y. H. Zhang et al., 2019). Moreover, because this gliotic response emerges concomitantly with the onset of recurrent seizures, seizure activity itself may further shape or amplify these glial changes (Vezzani et al., 2022), making it difficult to disentangle the primary disease mechanisms from secondary seizure-induced effects. Thus, while gliosis provides important insight into disease progression, it may be less well suited for dissecting the primary molecular mechanisms underlying the Arg87Cys gain-of-function. In contrast, we also observed that layer dispersion of hippocampal CA1 pyramidal neurons in *Cyfip2^+/R87C^* mice began to emerge before PWK 7 (Kang et al., 2023; Ma, Kang, et al., 2025). At this earlier stage, there is no overt neuronal loss or severe gliosis, making it a more suitable window to investigate neuron-intrinsic alterations and the molecular mechanisms underlying the Arg87Cys variant. We therefore asked whether, in contrast to CYFIP2 loss, which reduces both WRC components and RBPs (Zhang et al., 2024), the Arg87Cys variant disrupts the balance between these two CYFIP2-associated pathways, and whether such an imbalance contributes to early neuronal phenotypes, including CA1 layer dispersion.

To address this question, we aimed to characterize the early neuronal phenotypes of *Cyfip2^+/R87C^* mice at PWK 7 and to identify the associated molecular alterations in the hippocampal CA1 region. We first performed a detailed analysis of neuronal layer dispersion in the hippocampal CA1 region and confirmed that this phenotype is observed exclusively in *Cyfip2^+/R87C^* mice, but not in *Cyfip2^+/-^* mice. We then conducted a series of integrative analyses on hippocampal tissue, including single-nucleus RNA sequencing (snRNA-seq), quantitative global proteomics, and phosphoproteomics, together with complementary validation experiments and electrophysiological and morphological analyses of CA1 pyramidal neurons. Through these approaches, we found that the *CYFIP2* Arg87Cys variant leads to an imbalance between two major groups of CYFIP2-interacting partners, the WRC and RBPs, characterized by an increase in RBPs and a decrease in WRC components. In addition, altered phosphorylation of synaptic proteins was accompanied by increased excitatory synaptic transmission and dendritic alterations, most prominently in the more severely dispersed deep-layer CA1 pyramidal neurons. Furthermore, our data raise the possibility of molecular links between the up-regulation of RBPs and the emergence of neuronal layer dispersion. Collectively, extending our previous phenotypic characterization (Ma, Kang, et al., 2025; Ma et al., 2026), our study delineates a toxic gain-of-function mechanism for the Arg87Cys variant and provides a framework for understanding disease pathogenesis, as well as for guiding the development of mechanism-based therapeutic strategies in the future.

## Results

### Neuronal layer dispersion in the hippocampal CA1 region of *Cyfip2^+/R87C^* mice

To confirm our previous findings and better characterize the age-dependent progression of neuronal layer dispersion in the hippocampal CA1 region of *Cyfip2^+/R87C^* mice, we measured neuronal layer thickness, visualized by anti-Neuronal Nuclei (NeuN, RNA binding fox-1 homolog 3 [RBFOX3]) immunostaining, at PWK 1 and 7 (Fig. 1A). At both stages, CA1 neuronal layer thickness was increased in *Cyfip2^+/R87C^* mice compared to wild-type (WT) littermates, with the difference becoming more pronounced at PWK 7 than at PWK 1. As an internal control, dentate gyrus (DG) neuronal layer thickness was comparable between genotypes at both stages.

**Figure 1.**
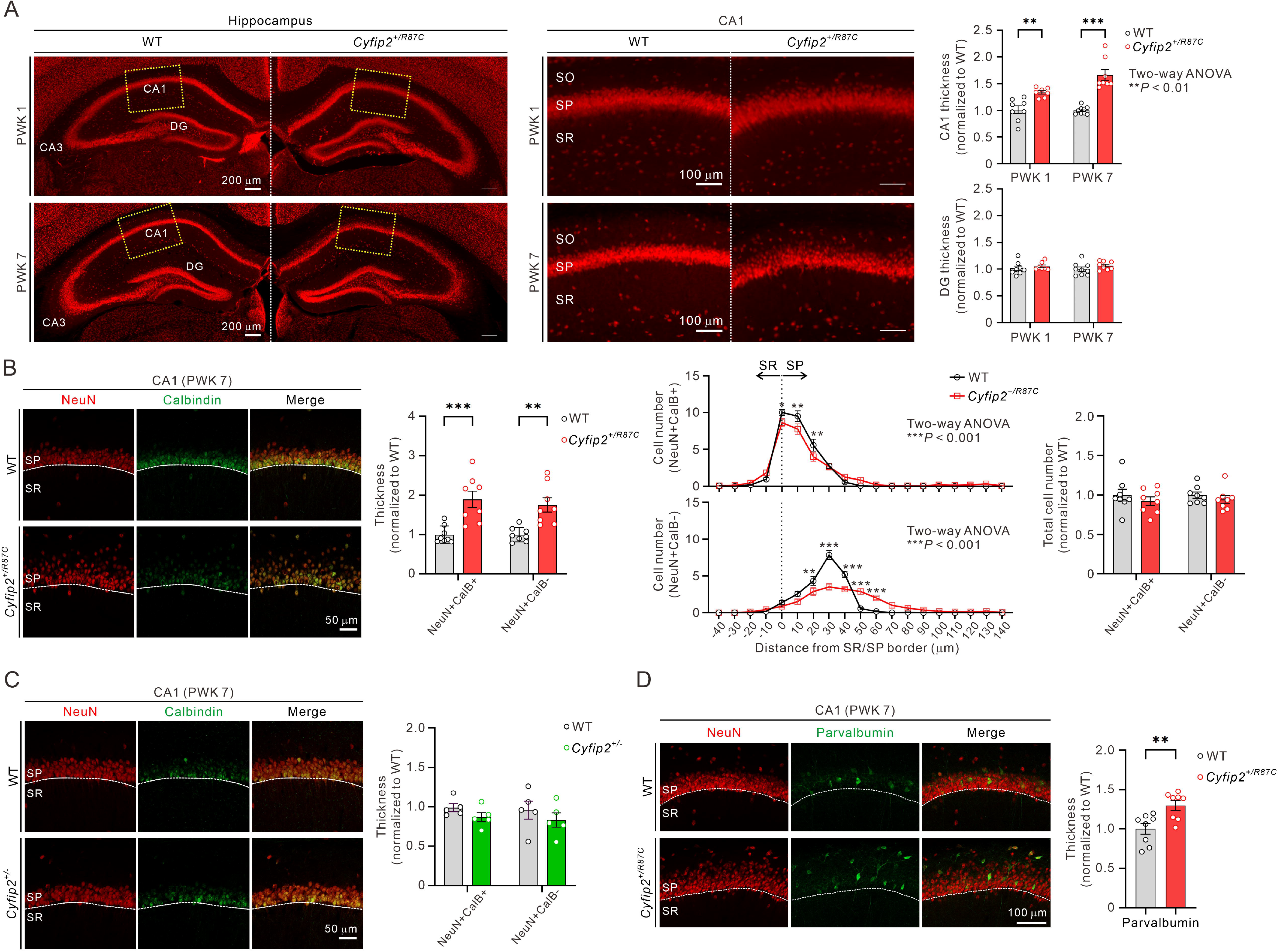
Neuronal layer dispersion in the hippocampal CA1 region of *Cyfip2^+/R87C^* mice. (**A**) Representative fluorescent immunohistochemistry images stained with anti-Neuronal Nuclei (NeuN) antibodies and corresponding quantification reveal neuronal layer dispersion in the Cornu Ammonis 1 (CA1), but not in the dentate gyrus (DG), of the hippocampus in *Cyfip2^+/R87C^* mice compared with wild-type (WT) mice at postnatal weeks 1 and 7 (PWK 1 and PWK 7) (n = 7–9 animals per genotype; two-way ANOVA with Šidák’s multiple-comparison test; the *P* value shown in the graph indicates a significant genotype × age interaction). The left panel shows the whole hippocampus, and the middle panel shows a magnified view of the CA1 region indicated by the yellow dashed box. SO, stratum oriens; SP, stratum pyramidale; SR, stratum radiatum. (**B**) Representative fluorescent immunohistochemistry images stained with anti-NeuN and anti-Calbindin (CalB) antibodies and corresponding quantification reveal neuronal dispersion in both superficial (CalB-positive) and deep (CalB-negative) layers of the hippocampal CA1 region in *Cyfip2^+/R87C^* mice (n = 8 animals per genotype; two-way ANOVA with Šidák’s multiple-comparison test; *P* values shown in the distribution graphs indicate significant genotype × distance interactions). Notably, analysis of cell number distribution along the radial axis, rather than layer thickness, suggests that dispersion is more pronounced in the deep layer. The total number of neurons in both superficial and deep layers does not differ significantly between *Cyfip2^+/R87C^* and WT mice. (**C**) Representative fluorescent immunohistochemistry images stained with anti-NeuN and anti-CalB antibodies and corresponding quantification reveal normal thickness of both superficial and deep layers of the hippocampal CA1 region observed in *Cyfip2^+/-^* mice at PWK 7 (n = 5 animals per genotype; two-way ANOVA with Šidák’s multiple-comparison test). (**D**) Representative fluorescent immunohistochemistry images stained with anti-NeuN and anti-parvalbumin antibodies and corresponding quantification reveal dispersion of parvalbumin-positive inhibitory interneurons in the hippocampal CA1 region of *Cyfip2^+/R87C^* mice at PWK 7 (n = 8 animals per genotype; unpaired two-tailed Welch’s t-test). Data are presented as mean ± SEM. \**P* < 0.05, \*\**P* < 0.01, \*\*\**P* < 0.001.

Hippocampal CA1 pyramidal neurons located in the stratum pyramidale are organized into two sublayers: a superficial layer (closer to stratum radiatum) and a deep layer (closer to stratum oriens), and anti-Calbindin (CalB) antibodies preferentially label superficial, but not deep, pyramidal neurons (Cid et al., 2021). Co-immunostaining for NeuN and CalB, together with measurement of cell layer thickness, revealed that both CalB-positive and CalB-negative neurons were more dispersed in *Cyfip2^+/R87C^* mice than in WT mice at PWK 7 (Fig. 1B). However, when we quantified neuronal distribution along the radial axis, CalB-negative (i.e., deep layer) pyramidal neurons exhibited larger changes in distribution than CalB-positive pyramidal neurons. The total numbers of CalB-positive and CalB-negative neurons were comparable between *Cyfip2^+/R87C^* mice and WT mice at PWK 7. This sublayer-specific difference prompted us to analyze superficial and deep pyramidal neurons separately in subsequent electrophysiological and morphological experiments (see Fig. 5).

In contrast, neither CalB-positive nor CalB-negative neuronal layer thickness was altered in *Cyfip2^+/-^* mice compared to WT littermates at PWK 7 (Fig. 1C). Intriguingly, not only pyramidal neurons but also parvalbumin-positive inhibitory interneurons, which also express CYFIP2 (S. H. Lee et al., 2020), showed significant dispersion in the CA1 region of *Cyfip2^+/R87C^* mice compared to WT mice (Fig. 1D). However, neuronal layer formation in the somatosensory cortex, assessed using layer-specific markers (brain-specific homeobox/POU domain protein 2 [Brn2], COUP-TF-interacting protein 2 [Ctip2], and cut-like homeobox 1 [Cux1]), was overall normal in *Cyfip2^+/R87C^* mice at PWK 7 (Supplementary Material, Fig. S1). Together, these findings indicate that the *Cyfip2* Arg87Cys variant, but not *Cyfip2* haploinsufficiency, causes a region-specific disruption of CA1 laminar organization, affecting both excitatory and inhibitory neurons.

### Exploratory snRNA-seq analysis nominates altered translation and cell adhesion/migration pathways in CA1 pyramidal neurons of *Cyfip2^+/R87C^* mice

To explore cell type-specific molecular changes in the hippocampus of *Cyfip2^+/R87C^* mice, we performed snRNA-seq on hippocampal tissue from WT and *Cyfip2^+/R87C^* mice at PWK 7. A total of 47,062 nuclei (26,060 from two WT and 21,002 from two *Cyfip2^+/R87C^* hippocampi) were analyzed. Clustering analysis, together with established marker gene expression and reference-based annotation using SingleR (Aran et al., 2019; Zhang et al., 2024), identified eleven distinct cell types (Fig. 2A and Supplementary Material, Fig. S2). Across both genotypes, excitatory neurons from the dentate gyrus (Excitatory-DG), excitatory CA1 pyramidal neurons (Excitatory-CA1), and oligodendrocytes represented the major cell populations. Comparison of cell type composition suggested a relative reduction in certain neuronal populations, most notably Excitatory-CA1 neurons and inhibitory interneurons, and a relative increase in glial populations (including astrocytes, microglia, oligodendrocytes, and oligodendrocyte precursor cells [OPCs]) in the *Cyfip2^+/R87C^* hippocampus compared with WT samples (Fig. 2B). However, given the limited number of biological replicates, these apparent differences were not subjected to formal statistical testing and were not interpreted further. Notably, histological quantification revealed no change in the total number of CA1 pyramidal neurons (Fig. 1B), suggesting that the apparent reduction in Excitatory-CA1 nuclei may reflect technical variability in tissue dissection or nuclei recovery rather than neuronal loss.

**Figure 2.**
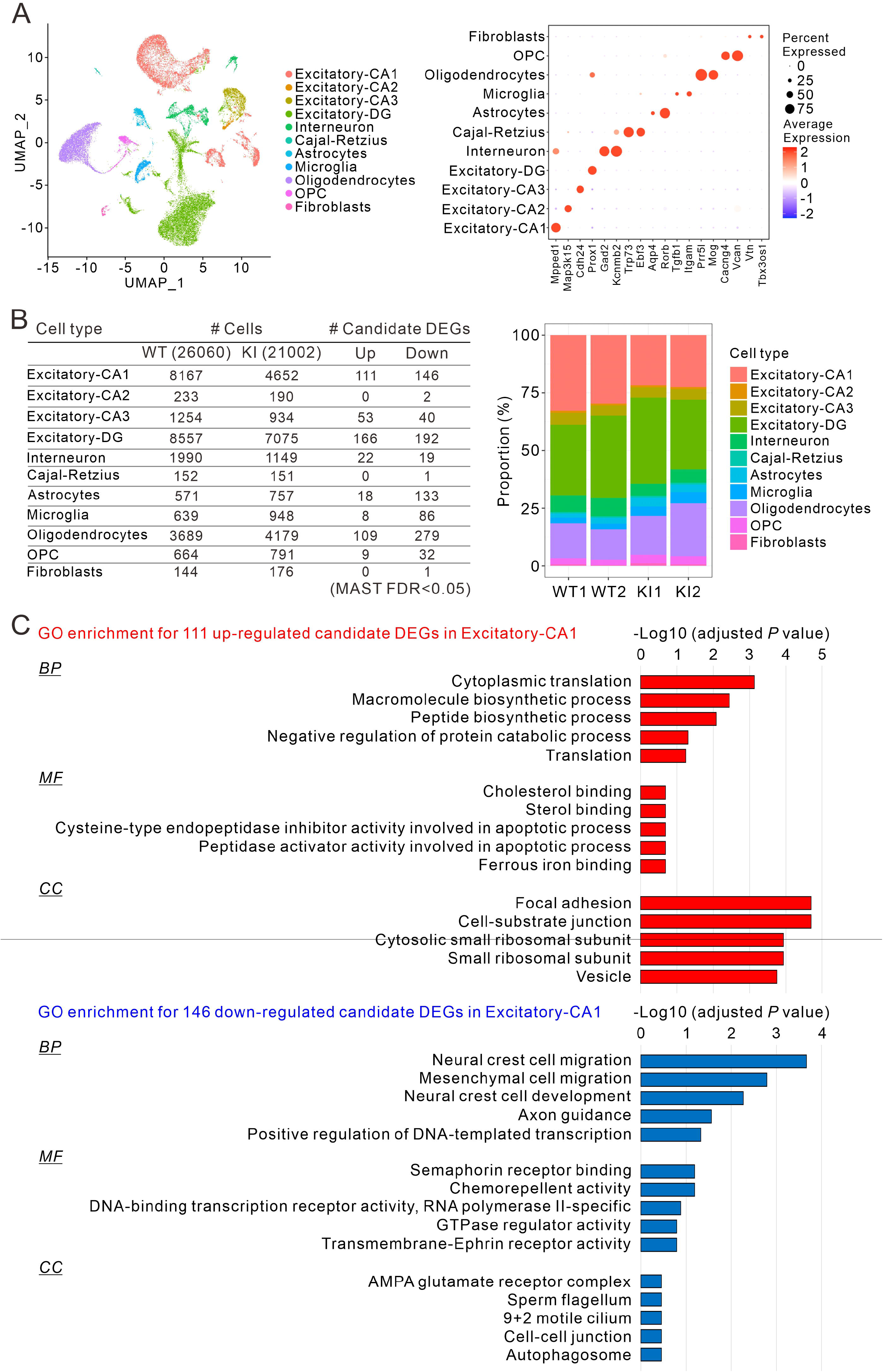
Exploratory single-nucleus transcriptomic analysis of the hippocampus in *Cyfip2^+/R87C^* mice. (**A**) Uniform manifold approximation and projection (UMAP) visualization of eleven cell types identified in the hippocampus at PWK 7 (left panel; nuclei from two WT and two *Cyfip2^+/R87C^* [KI] mice were integrated). The right panel displays a dot plot illustrating the expression of representative marker genes for each cell type; dot size indicates the percentage of nuclei expressing each gene, and color indicates scaled average expression. OPC, oligodendrocyte precursor cell. (**B**) Summary of cell numbers and candidate differentially expressed genes (DEGs; MAST, false discovery rate [FDR] < 0.05) across the eleven cell types (left panel). The right panel shows the proportions of the eleven cell types in each of the two WT and two *Cyfip2^+/R87C^* hippocampal samples; given the limited number of biological replicates, these proportions were not statistically tested. (**C**) Bar graphs showing the results of Gene Ontology over-representation analysis (Enrichr) for the 111 candidate upregulated (red) and 146 candidate downregulated (blue) DEGs in Excitatory-CA1 neurons. For each Gene Ontology category, the five top-ranked terms are shown, irrespective of statistical significance, for exploratory purposes; *P* values were adjusted using the Benjamini-Hochberg method. Terms with adjusted *P* < 0.05 are listed in Supplementary Material, Table S2. BP, biological process; CC, cellular component; MF, molecular function.

Using this dataset as an exploratory resource, we next examined candidate transcriptional changes across the annotated cell populations. The largest numbers of candidate expression changes were observed in oligodendrocyte, Excitatory-DG, and Excitatory-CA1 populations, whereas relatively few were detected in Excitatory-CA2, Cajal-Retzius, and fibroblast cell types, potentially reflecting differences in cell abundance (Fig. 2B and Supplementary Material, Table S1). To generate hypotheses regarding pathways potentially linked to the CA1 dispersion phenotype, we performed Gene Ontology (GO) over-representation analysis using candidate expression changes in the Excitatory-CA1 population. The candidate upregulated gene set suggested enrichment of pathways related to translation, ribosome function, and focal adhesion, whereas the candidate downregulated gene set suggested enrichment of pathways associated with cell migration, axon guidance, including semaphoring and ephrin receptor genes (Fig. 2C). Other cell types showed distinct enrichment patterns (Supplementary Material, Table S2). Collectively, these exploratory snRNA-seq analyses generated cell type-specific hypotheses, particularly implicating altered translation and cell adhesion/migration-related pathways in CA1 pyramidal neurons.

### HEK293T cells expressing CYFIP2 Arg87Cys exhibit reduced proliferation, morphological alterations, and changes in cell adhesion–related gene expression

Although our immunostaining and snRNA-seq analyses revealed cell type-specific alterations in the hippocampus of *Cyfip2^+/R87C^* mice, we reasoned that, given the cellular heterogeneity and potential crosstalk in this region, further experiments were required to determine whether the *CYFIP2* Arg87Cys variant exerts cell-autonomous effects. We therefore turned to a simpler and more tractable system using human embryonic kidney 293T (HEK293T) cells. Using the Clustered Regularly Interspaced Short Palindromic Repeats (CRISPR)/CRISPR-associated protein 9 (Cas9) system, we generated two independent clonal lines each of heterozygous (+/R87C) and homozygous (R87C/R87C) *CYFIP2* Arg87Cys mutant HEK293T cells, in which one or both alleles were edited, respectively (Fig. 3A). We detected no editing at nine predicted off-target sites in these clones (Fig. 3A). For all subsequent experiments, one representative clonal line for each genotype was used.

**Figure 3.**
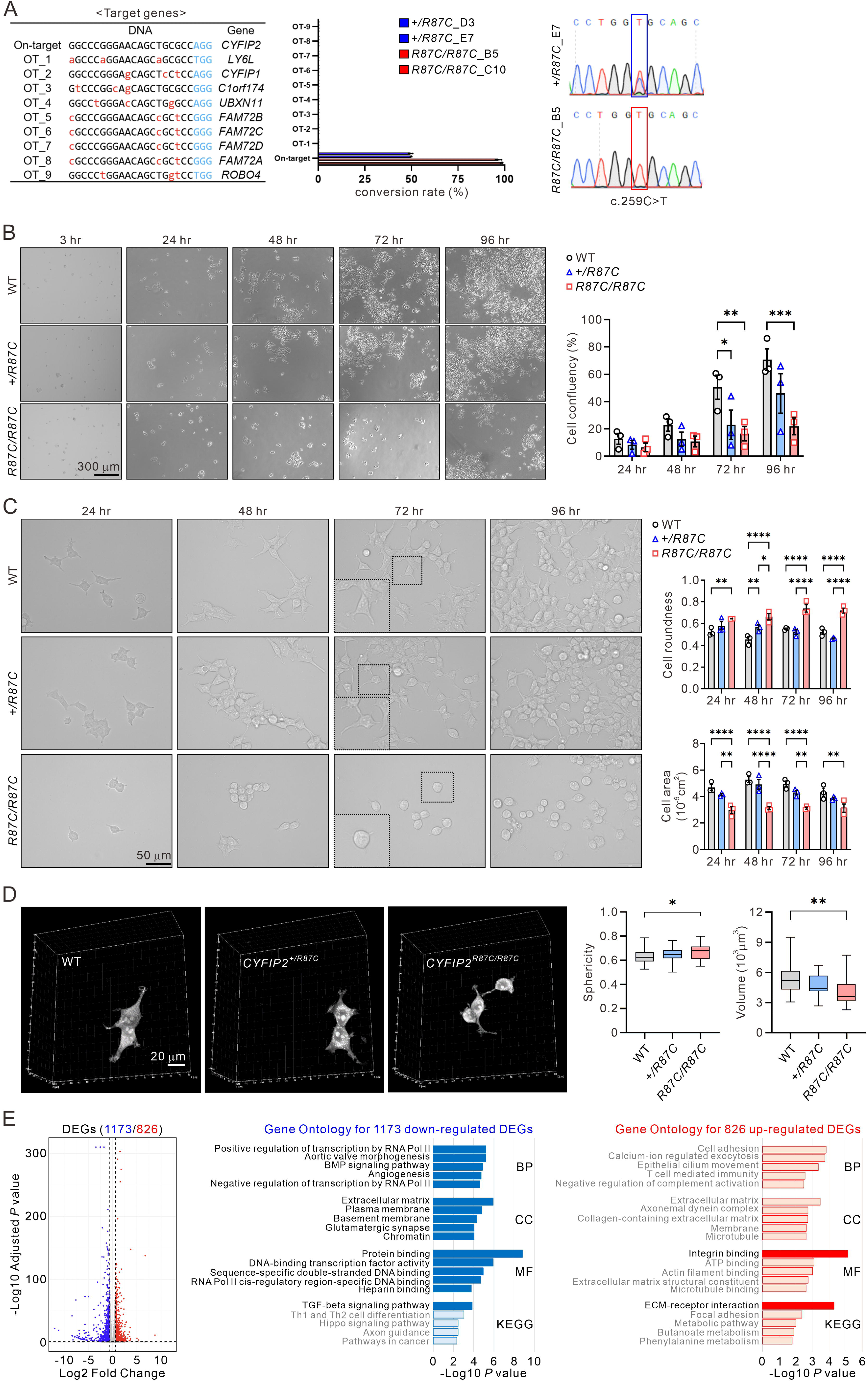
Generation and characterization of HEK293T cells expressing CYFIP2 Arg87Cys. (**A**) List of on-and off-target sites used for gene editing, together with targeted deep sequencing results showing conversion rates at these sites in two heterozygous (+/R87C_D3 and +/R87C_E7) and two homozygous (R87C/R87C_B5 and R87C/R87C_C10) cell lines. Red lowercase letters indicate mismatches relative to the on-target sequence, and blue letters indicate protospacer adjacent motif (PAM) sequences. Conversion was detected only at the on-target site, with no editing at any of the nine predicted off-target sites (OT_1–OT_9). Representative Sanger sequencing chromatograms showing the target sequence change (*CYFIP2* c.259C>T) in the heterozygous and homozygous cell lines. (**B**) Representative 2D brightfield images of WT, +/R87C, and R87C/R87C HEK293T cells acquired at various time points after seeding onto culture dishes. A graph depicting the confluency of the cell lines over time (n = 3 independent experiments using separate cell culture preparations from different passages; two-way ANOVA with Tukey’s post hoc analysis). (**C**) Representative 2D brightfield images of WT, +/R87C, and R87C/R87C HEK293T cells, together with quantification of cell roundness and area at different time points post-seeding (n = 3 independent experiments using separate cell culture preparations from different passages; two-way ANOVA with Tukey’s post hoc analysis). (**D**) Representative 3D holotomography images of WT, +/R87C, and R87C/R87C HEK293T cells, together with quantification of sphericity and volume (n = 3 independent experiments using separate cell culture preparations from different passages, with 21–33 cells analyzed per genotype; one-way ANOVA with Tukey’s post hoc analysis). Box plots show the median (center line), interquartile range (box; 25th–75th percentiles), and minimum to maximum values (whiskers). (**E**) Volcano plot of differentially expressed genes (DEGs) identified in R87C/R87C HEK293T cells compared with WT cells (blue, 1,173 down-regulated; red, 826 up-regulated). Dashed lines indicated the thresholds used to define DEGs (adjusted *P* value = 0.05 and absolute fold change = 1.5). Bar graphs show enriched terms from Gene Ontology (GO) and Kyoto Encyclopedia of Genes and Genomes (KEGG) pathway analyses for down-regulated (blue) and up-regulated (red) DEGs, with the five top-ranked terms shown for each category. Light-colored bars indicate non-significant terms. Data are presented as mean ± SEM. \**P* < 0.05, \*\**P* < 0.01, \*\*\**P* < 0.001, \*\*\*\**P* < 0.0001.

The most prominent phenotype observed in these *CYFIP2* Arg87Cys mutant cell lines was a marked reduction in cell growth, as assessed by confluency over time (Fig. 3B). Confluency was reduced in both +/R87C and R87C/R87C cells compared with WT cells at 72 h, and remained significantly reduced in R87C/R87C cells at 96h, although the two mutant genotypes did not differ significantly from each other. A second characteristic was a change in cell morphology. Mutant cells exhibited increased roundness and reduced cell size (Fig. 3C), which was further confirmed by three-dimensional analyses using holotomography imaging (Fig. 3D). For these morphological measures, R87C/R87C cells differ significantly not only from WT cells but also from +/R87C cells at several time points, consistent with an effect that scales with Arg87Cys allele dosage.

To gain molecular insights into these cellular phenotypes, we performed RNA-seq analysis on WT and R87C/R87C HEK293T cells (Fig. 3E). Gene Ontology (GO) and Kyoto Encyclopedia of Genes and Genomes (KEGG) pathway analyses of DEGs (1173 down-regulated and 826 up-regulated, Supplementary Material, Table S3) revealed a down-regulation of RNA polymerase II and transcription-related processes, together with enrichment of extracellular matrix-and membrane-related terms among the down-regulated DEGs. Among the up-regulated DEGs, the most significantly enriched terms were integrin binding and ECM-receptor interaction, with additional adhesion-related terms such as cell adhesion and focal adhesion showing a similar tendency. Given that cell adhesion, morphology, and proliferation are highly interconnected processes (Jones et al., 2019), these RNA-seq results are consistent with the observed cytological changes in *CYFIP2* Arg87Cys mutant HEK293T cells. Although HEK293T cells are proliferative and differ substantially from neurons, these results raise the possibility that the *CYFIP2* Arg87Cys variant may exert cell-autonomous effects on pathways related to cell adhesion.

### Quantitative global proteomic analysis reveals up-regulation of RBPs and down-regulation of WRC components in the CA1 region of *Cyfip2^+/R87C^* mice

Next, we examined the impact of the *CYFIP2* Arg87Cys variant at the protein level, as CYFIP2 is not a transcription factor, and thus changes at the proteomic level may more directly capture molecular mechanisms underlying the cellular and transcriptomic phenotypes described above. To this end, we performed quantitative global proteomic analysis on microdissected hippocampal CA1 regions from eight pairs of *Cyfip2^+/R87C^* and WT mice at PWK 7 (Fig. 4A). Among a total of 6,773 detected proteins, we identified 1,334 significantly altered differentially expressed proteins (DEPs) (Supplementary Material, Table S4). Of these, 693 proteins were up-regulated and 641 were down-regulated in *Cyfip2^+/R87C^* mice compared with WT controls (Fig. 4B). Notably, CYFIP2 and other WRC components (ABI1, ABI2, NCKAP1, and WAVE1) were among the most prominently down-regulated DEPs. This finding is consistent with the reduced stability of the CYFIP2 protein caused by the Arg87Cys variant (Kang et al., 2023; Lee et al., 2019), as well as the interdependent protein stability among WRC components (Zhang et al., 2018; Zhang et al., 2020), and thus supports the robustness and validity of our proteomic analysis.

**Figure 4.**
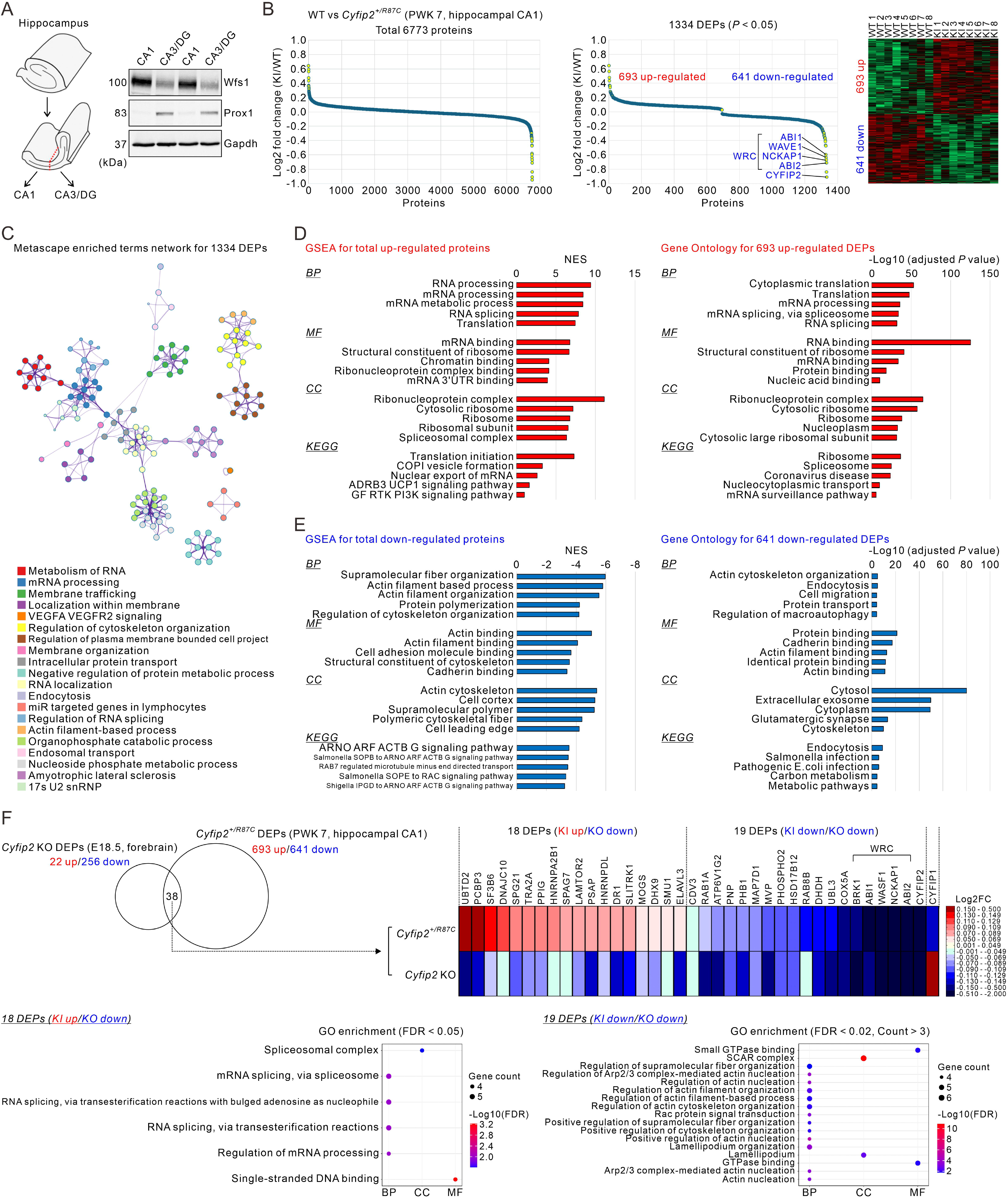
Quantitative global proteomic analysis of the hippocampal CA1 region in *Cyfip2^+/R87C^* mice. (**A**) A schematic diagram illustrates microdissection of the CA1 region and the remaining hippocampal tissue (i.e., CA3/DG). Representative Western blot images validate the accuracy of the microdissection (two representative samples per region are shown). Wolfram syndrome 1 (Wfs1) and prospero homeobox 1 (Prox1) are markers for CA1 pyramidal neurons and DG granule cells, respectively. Glyceraldehyde-3-phosphate dehydrogenase (Gapdh) is used as a loading control. (**B**) Relative protein abundance (total 6,773 identified proteins) comparison between the hippocampal CA1 regions of *Cyfip2^+/R87C^*knock-in (KI) and wild-type (WT) mice at PWK 7 (left panel). The middle panel shows the relative abundance of 1,334 DEPs (693 up-regulated and 641 down-regulated) between *Cyfip2^+/R87C^* KI and WT mice. Components of the WAVE regulatory complex (WRC) and CYFIP2 are indicated on the plot. The right panel presents a heatmap of the 1,334 DEPs for eight pairs of *Cyfip2^+/R87C^* KI and WT samples; red and green indicate higher and lower relative abundance, respectively. (**C**) A network map of Metascape enriched terms for the 1,334 DEPs. Each node represents an enriched term, nodes are colored by cluster, and edges connect terms with similarity above the Metascape default threshold. (**D**) Bar graph showing the results of gene set enrichment analysis (GSEA) for the total up-regulated proteins, regardless of statistical significance (left panel; NES, normalized enrichment score). Bar graph displaying the results of GO and KEGG pathway analyses for the 693 up-regulated DEPs (right panel). (**E**) Bar graph showing the results of GSEA for the total down-regulated proteins, regardless of statistical significance (left panel). Bar graph displaying the results of GO and KEGG pathway analyses for the 641 down-regulated DEPs (right panel). (**F**) A Venn diagram illustrates the 38 proteins shared between the 278 DEPs identified in the *Cyfip2* knock-out (KO) forebrain at embryonic day 18.5 (E18.5) and the 1,334 DEPs identified in *Cyfip2^+/R87C^* KI mice. The heatmap depicts expression changes of these 38 shared DEPs in *Cyfip2* KO and *Cyfip2^+/R87C^* mice relative to their respective WT controls. Among them, 18 DEPs are up-regulated in *Cyfip2^+/R87C^* mice but down-regulated in *Cyfip2* KO mice, whereas 19 DEPs are commonly down-regulated in both mouse models. In contrast, CYFIP1 is down-regulated in *Cyfip2^+/R87C^* mice but up-regulated in *Cyfip2* KO mice. Over-representation analyses were performed for the 18 KI up-/KO down-regulated DEPs (bottom left) and the 19 KI down-/KO down-regulated DEPs (bottom right). FC, fold change; FDR, false discovery rate.

Network analysis of enriched terms using Metascape of the entire set of 1,334 DEPs revealed significant enrichment of terms related to RNA metabolism, RNA processing and localization, as well as actin cytoskeleton organization (Fig. 4C). These results are similar to those from our previous quantitative global proteomic analysis of embryonic *Cyfip2*-null mouse forebrains (Zhang et al., 2024). However, in the *Cyfip2*-null forebrain, both RBPs and WRC components were down-regulated, with approximately 92% of all DEPs (256 out of 278) showing decreased expression. In contrast, in *Cyfip2^+/R87C^* mice, up-regulated and down-regulated DEPs were identified in nearly equal proportions. We therefore next examined whether the up-regulated and down-regulated DEPs in *Cyfip2^+/R87C^* mice were associated with distinct biological pathways.

Consistent with this, GSEA of the total set of up-regulated proteins (regardless of statistical significance), together with GO and KEGG pathway analyses of significantly up-regulated DEPs in *Cyfip2^+/R87C^* mice, commonly revealed significant enrichment of terms related to RNA processing, splicing, translation, and ribosome (Fig. 4D and Supplementary Material, Table S5 and S6). In contrast, the same analyses applied to the down-regulated proteins indicated enrichment for pathways associated with actin cytoskeleton organization and binding (Fig. 4E and Supplementary Material, Table S7 and S8). Together, these results suggest that, unlike the embryonic *Cyfip2*-null forebrain, the hippocampal CA1 region of *Cyfip2^+/R87C^* mice at PWK 7 exhibits a distinct global proteomic pattern characterized by an increase in RBPs and a decrease in WRC components. Although the two analyses were performed on brain samples that differ in developmental stage (embryonic vs. PWK 7) and anatomical region (forebrain vs. hippocampal CA1), the opposite directions of RBP regulation observed in the two models are unlikely to be explained solely by these differences and instead point to fundamentally different molecular consequences of CYFIP2 loss versus the Arg87Cys variant.

To more directly assess these differences, we compared the 278 DEPs identified in *Cyfip2*-null mice with the 1,334 DEPs from *Cyfip2^+/R87C^* mice (Fig. 4F). A total of 38 proteins were commonly detected between the two datasets. Among these, 18 proteins were up-regulated in *Cyfip2^+/R87C^* mice but down-regulated in *Cyfip2*-null mice, whereas 19 proteins were down-regulated in both models. Only CYFIP1 showed an opposite pattern, being decreased in *Cyfip2^+/R87C^* mice but increased in *Cyfip2*-null mice. Over-representation analysis of these protein groups revealed that the 18 oppositely regulated proteins (excluding CYFIP1) were significantly enriched for terms related to RNA processing and splicing, while the 19 commonly down-regulated proteins were enriched for actin nucleation and cytoskeleton organization, consistent with their inclusion of WRC components. Together, these results are consistent with the interpretation described above.

### Quantitative phosphoproteomic analysis reveals altered synaptic protein phosphorylation accompanied by functional and dendritic alterations in CA1 pyramidal neurons of *Cyfip2^+/R87C^* mice

Following the quantitative global proteomic analysis, we performed quantitative phosphoproteomic analysis on the same CA1 samples. This approach was intended to identify proteins with altered activity states, which can often be more sensitively captured by changes in phosphorylation than by changes in total protein abundance, and thereby providing deeper insights into the neuronal alterations, potentially beyond layer dispersion, in *Cyfip2^+/R87C^*mice.

In total, 11,845 phosphopeptides were identified across WT and *Cyfip2^+/^*^R87C^ samples (Fig. 5A and Supplementary Material, Table S9). To identify phosphopeptides specifically altered in *Cyfip2^+/R87C^* mice relative to WT, normalization using the results of the global proteomic analysis was required. This step was performed to exclude phosphorylation changes that merely reflected corresponding alterations in total protein abundance (Fig. 5A and see methods for details). When the retained phosphopeptides (|Z| > 1.96, *P* < 0.05) were mapped back to their corresponding proteins (i.e., multiple phosphopeptides derived from the same protein were considered as a single protein), a total of 344 proteins were identified. Among these, 252 proteins exhibited increased phosphorylation, whereas 118 proteins showed decreased phosphorylation in *Cyfip2^+/R87C^* mice compared with WT mice. Notably, for 26 proteins, both increased and decreased phosphorylation were detected, depending on the specific phosphosite. Among the proteins showing the largest increases in phosphorylation fold change in *Cyfip2^+/R87C^* mice were synaptic Ras GTPase-activating protein 1 (SynGAP1), synapsin I (Syn1), microtubule affinity-regulating kinase 4 (Mark4), microtubule-associated protein 2 (Map2), and diacylglycerol kinase beta (Dgkβ). In contrast, proteins exhibiting the largest decreases in phosphorylation fold change included protein kinase C epsilon (Prkcε), bassoon presynaptic cytomatrix protein (Bsn), phosphodiesterase 4A (Pde4A), and ribosomal protein lateral stalk subunit P2 (Rplp2).

**Figure 5.**
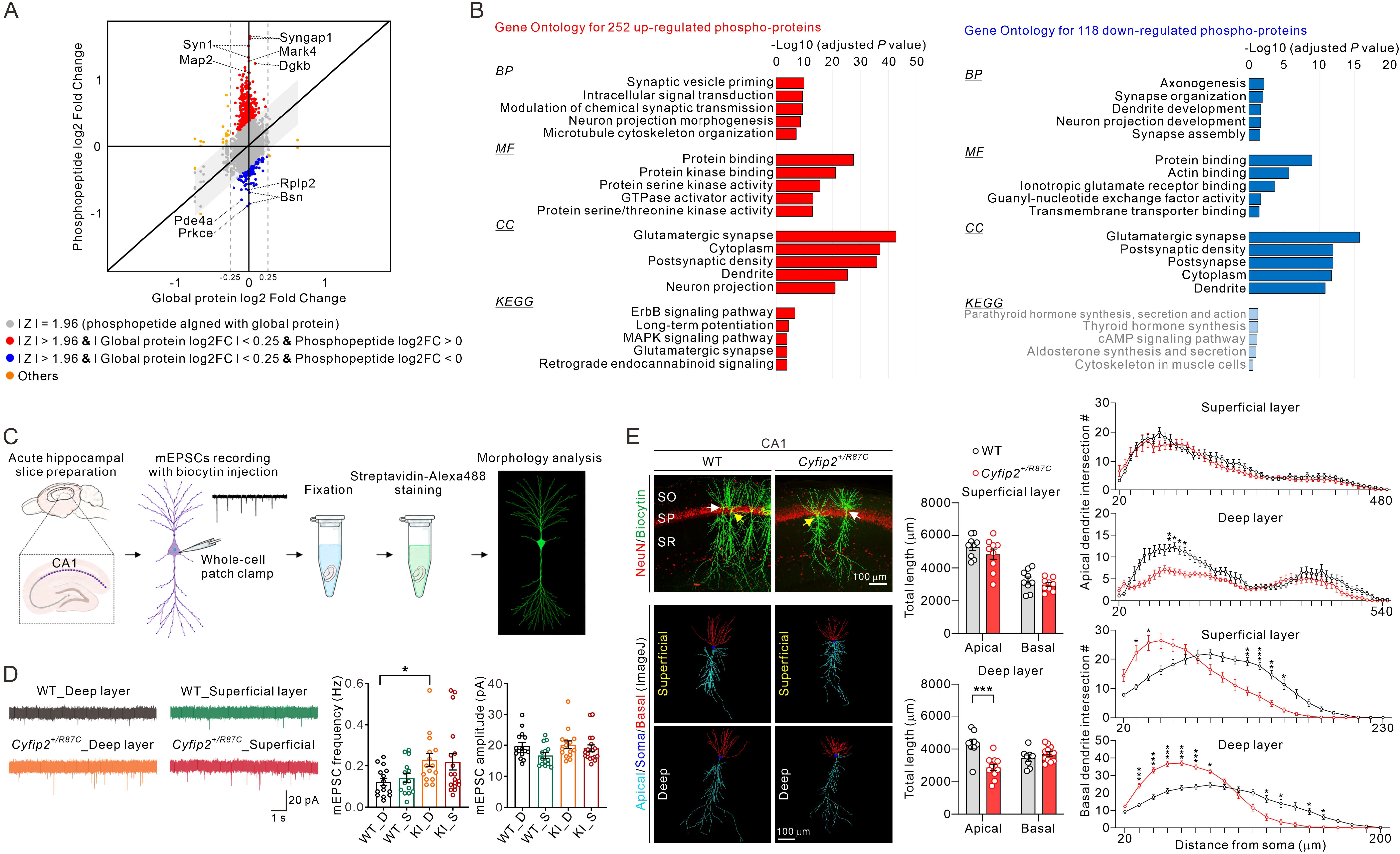
Quantitative phosphoproteomic analysis and synaptic and dendritic characterization of pyramidal neurons in the hippocampal CA1 region in *Cyfip2^+/R87C^* mice. (**A**) Scatter plot illustrating the relationship between phosphopeptide log fold change (FC) and the corresponding global protein log FC. Each point represents an individual phosphopeptide. Phosphopeptides were classified based on diagonal-based z-score analysis, and nonsignificant peptides (gray, |z| ≤ 1.96) were excluded from emphasis. Among significant phosphopeptides (|z| > 1.96), those with minimal protein-level change (|Global protein log FC| < 0.25) were colored red or blue depending on the direction of deviation from the diagonal (red: Phospho log FC > 0; blue: Phospho log FC < 0), highlighting phosphorylation-specific modulation. Phosphopeptides that significantly deviated but were associated with a substantial protein-level change (|Global protein log FC| ≥ 0.25) were colored orange, suggesting compensatory phosphorylation responses that may buffer or counteract broader protein expression changes. Representative proteins showing the largest increase or decreases in phosphorylation are indicated. (**B**) Bar graphs show enriched terms from GO and KEGG pathway analyses for the 252 up-regulated (red) and 118 down-regulated (blue) phospho-proteins; the five top-ranked terms are shown for each category. Light-colored bars indicate non-significant terms. (**C**) Schematic of the experimental workflow. Whole-cell patch-clamp recordings of miniature excitatory postsynaptic currents (mEPSCs) were obtained from CA1 pyramidal neurons in acute hippocampal slices from WT and *Cyfip2^+/R87C^* mice at PWK 6–9, with biocytin included in the pipette solution for post hoc visualization with Streptavidin-Alexa 488 and subsequent morphological analysis. (**D**) Representative mEPSC traces and quantification of mEPSC frequency an amplitude in superficial-layer (S) an deep-layer (D) CA1 pyramidal neurons of WT and *Cyfip2^+/R87C^* knock-in (KI) mice (n = 14–18 neurons from 6 animals per genotype; two-way ANOVA with Šidák’s multiple-comparison test). (**E**) Representative images of biocytin-filled CA1 pyramidal neurons counterstained with anti-NeuN antibodies (top left); yellow and white arrows indicate superficial- and deep-layer neurons, respectively. Representative ImageJ reconstructions of superficial- and deep-layer neurons are shown below, with apical dendrites in cyan, basal dendrites in red, and the soma in blue. Quantification of total apical and basal dendritic length (middle) and Sholl analysis of apical and basal dendritic (right) are shown separately for superficial- and deep-layer neurons (n = 14–18 neurons from 6 animals per genotype; two-way ANOVA with Šidák’s multiple-comparison test). Data are presented as mean ± SEM. \**P* < 0.05, \*\**P* < 0.01, \*\*\**P* < 0.001.

To characterize the functional properties of these proteins, we performed GO and KEGG pathway analyses of the 252 up-regulated and 118 down-regulated phosphoproteins. Notably, synapse-related pathway terms, including glutamatergic synapse, postsynaptic density, and synaptic transmission, were significantly enriched in both groups (Fig. 5B and Supplementary Material, Table S10 and S11). These results suggest alterations in the phosphorylation states of glutamatergic excitatory synaptic proteins in hippocampal CA1 neurons of *Cyfip2^+/R87C^* mice at PWK 7.

To determine whether these molecular changes are associated with functional alterations in excitatory synaptic transmission, we next performed whole-cell patch-clamp recordings of miniature excitatory postsynaptic currents (mEPSCs) from CA1 pyramidal neurons in acute hippocampal slices from WT and *Cyfip2^+/R87C^* mice at PWK 6–9 (Fig. 5C). Given the layer dispersion observed in the CA1 region of *Cyfip2^+/R87C^* mice, pyramidal neurons were analyzed separately according to their laminar position. In WT mice, recordings were obtained from neurons located in the superficial and deep layers of the CA1 pyramidal cell layer. Corresponding superficial-and deep-layer neurons were examined in *Cyfip2^+/R87C^* mice, with the latter including dispersed deep-layer neurons. During whole-cell recording, neurons were filled with biocytin for subsequent visualization and morphological analysis following fixation and staining with Streptavidin-Alexa 488 (Fig. 5C).

Analysis of mEPSCs revealed an overall tendency toward increased event frequency in CA1 pyramidal neurons from *Cyfip2^+/R87C^* mice, with a significant increase observed in deep-layer neurons compared with WT deep-layer neurons (Fig. 5D). In contrast, mEPSC amplitudes were generally comparable between genotypes and across laminar populations (Fig. 5D). These findings indicate altered excitatory synaptic transmission in *Cyfip2^+/R87C^*CA1 pyramidal neurons, particularly in the deep-layer population, characterized by increased mEPSC frequency without a corresponding change in mEPSC amplitude. This pattern, together with the altered phosphorylation of multiple presynaptic proteins, such as Syn1 and Bsn, raises the possibility of altered presynaptic function in *Cyfip2^+/R87C^* CA1 neurons.

To directly examine whether these phosphoproteomic changes are accompanied by structural alterations in excitatory synapses, we performed fluorescent immunohistochemistry on hippocampal sections from WT and *Cyfip2^+/R87C^* mice at PWK 7 using antibodies against vesicular glutamate transporter 1 (vGluT1) and Homer1, which label excitatory presynaptic and postsynaptic compartments, respectively (Supplementary Material, Fig. S3). However, detailed quantitative analyses revealed no significant differences between *Cyfip2^+/R87C^* and WT mice in puncta intensity, volume, density, or colocalization in dendritic regions of CA1 pyramidal neurons. These findings are consistent with our previous report showing no significant changes in the expression levels of representative pre- and postsynaptic proteins in the hippocampus of *Cyfip2^+/R87C^* mice at PWK 7 (Ma, Kang, et al., 2025).

We next analyzed the morphology of the biocytin-filled CA1 pyramidal neurons from which mEPSCs had been recorded, reconstructing apical and basal dendrites separately in superficial- and deep-layer neurons (Fig. 5E). In superficial-layer neurons, the total lengths of both apical and basal dendrites were comparable between *Cyfip2^+/R87C^* and WT mice. In deep-layer neurons, however, the total apical dendritic length was significantly reduced in *Cyfip2^+/R87C^* mice, whereas the total basal dendritic length remained unchanged (Fig. 5E). Sholl analysis revealed that dendritic branching patterns were altered even when total dendritic length was preserved (Fig. 5E). Apical dendrites of superficial-layer neurons showed comparable intersection profiles between genotypes, whereas those of deep-layer neurons exhibited significantly fewer intersections at proximal to intermediate distances from the soma in *Cyfip2^+/R87C^* mice, consistent with their reduced total apical length. In contrast, basal dendrites displayed a redistribution of branching in both sublayers: intersections were increased at proximal distances and decreased at distal distances in *Cyfip2^+/R87C^* mice compared to WT mice. This redistribution was more pronounced in deep-layer neurons, in which the proximal increase was markedly larger than those observed in superficial-layer neurons.

Together these results indicate that CA1 pyramidal neurons of *Cyfip2^+/R87C^* mice at PWK 7 exhibit altered excitatory synaptic transmission and dendritic architecture, in line with the altered phosphorylation states of synaptic and cytoskeletal proteins described above. Notably, both the increase in mEPSC frequency and the dendritic alterations were most prominent in deep-layer neurons, which also show more severe dispersion than superficial-layer neurons (Fig. 1B). These observations suggest that the synaptic and dendritic changes are closely associated with the altered laminar positioning of CA1 pyramidal neurons in *Cyfip2^+/R87C^*mice.

### EphA3 down-regulation as a candidate link between RBP up-regulation and neuronal layer dispersion

Based on previous studies (Y. Lee et al., 2020; Zhang et al., 2024) together with our new data, we confirmed that CYFIP2 physically and functionally interacts with two distinct groups of proteins: the WRC and associated actin regulators, and RBPs (Fig. 6A, panel 1). In the absence of CYFIP2, both groups are concomitantly reduced (Fig. 6A, panel 2). In contrast, expression of CYFIP2 Arg87Cys produces a divergent pattern, with decreased WRC components accompanied by increased RBPs, resulting in an imbalance between these two CYFIP2-associated functional modules (Fig. 6A, panel 3). Although we cannot exclude the possibility that CYFIP2 Arg87Cys may also engage additional, previously unrecognized protein partners, we sought to explore a potential molecular link between two features uniquely observed in *Cyfip2^+/R87C^* mice, RBP up-regulation and CA1 neuronal layer dispersion, possibly involving pathways related to cell adhesion and/or migration. One plausible scenario is that increased RBPs alter the translation of their target mRNAs, particularly those encoding proteins involved in cell adhesion and/or migration.

**Figure 6.**
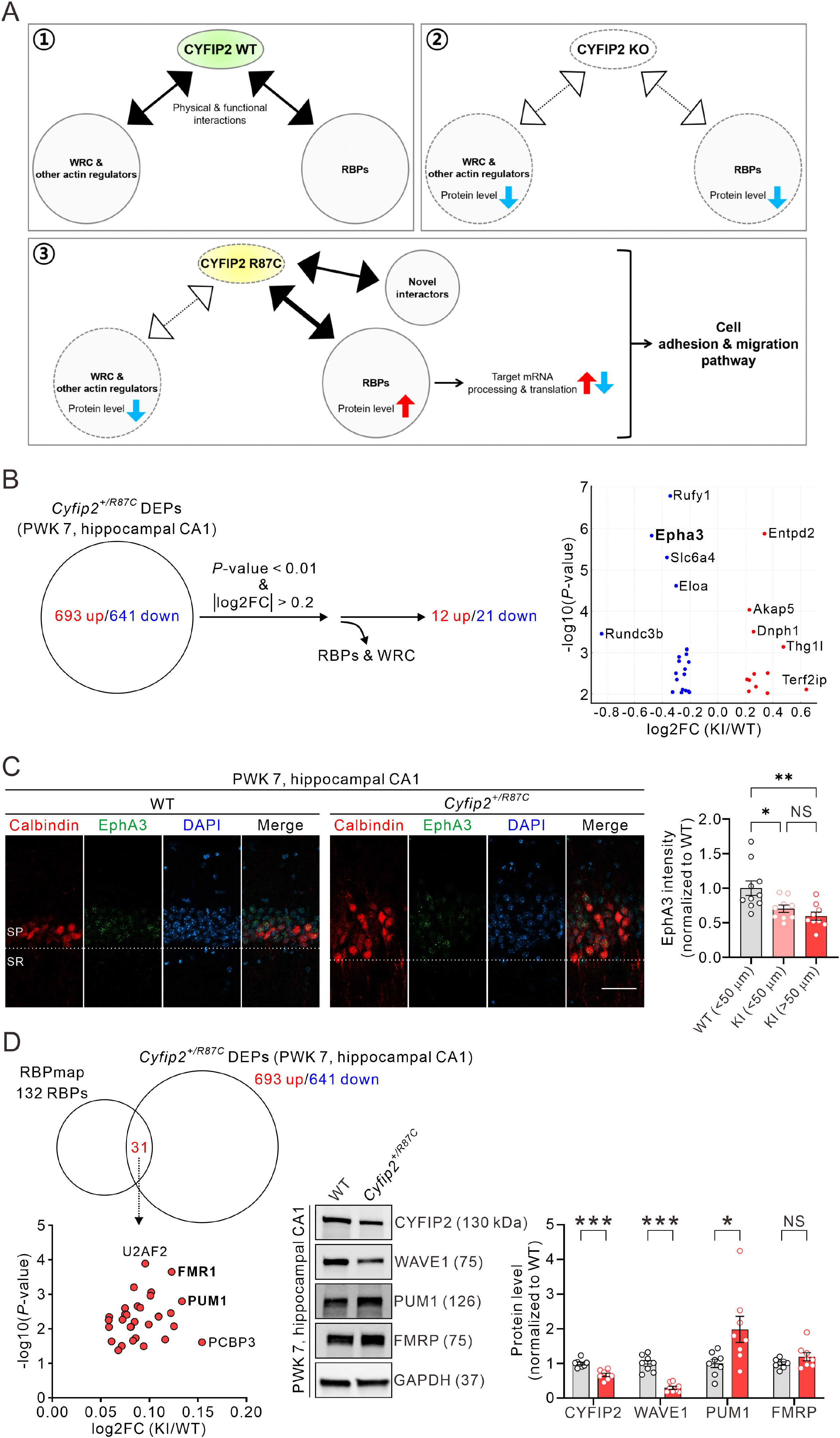
Potential involvement of EphA3 down-regulation in RBP-associated neuronal layer dispersion. (**A**) Schematic diagrams illustrating the dual molecular functions and interactions of CYFIP2 (i.e., WRC and RBPs; panel 1), and their potential alterations upon loss of CYFIP2 (CYFIP2 KO; panel 2) or the Arg87Cys variant; panel 3. In the Arg87Cys condition, CYFIP2 is proposed to exhibit reduced association with the WRC and enhanced interactions with RBPs and/or potential novel interactors. This shift may contribute to increased RBP-mediated regulation of target mRNA processing and translation. Collectively, these molecular changes are hypothesized to impact cell adhesion and migration pathways. Blue and red arrows indicate decreased and increased protein levels, respectively. (**B**) Summary of the process used to identify candidate proteins among the DEPs potentially affected by RBP up-regulation in *Cyfip2^+/R87C^* mice. From the 1,334 DEPs, proteins with the criteria of *P* < 0.01 and |log2 fold change| > 0.2 were selected. WRC components and RBPs were then excluded, resulting in 12 up-regulated and 21 down-regulated proteins. The graph on the right shows the *P* values and fold changes of these 33 proteins; EphA3 is highlighted in bold. (**C**) Representative fluorescent immunohistochemistry images stained with anti-CalB and anti-EphA3 antibodies, with nuclei counterstained with 4′,6-diamidino-2-phenylindole (DAPI), and corresponding quantification show reduced EphA3 expression in CalB-positive neurons in the hippocampal CA1 region of *Cyfip2^+/R87C^* mice at PWK 7. CalB-positive neurons were grouped according to their distance from the boundary between the stratum pyramidale (SP) and stratum radiatum (SR), indicated by the dotted line (n = 9 to 10 animals per genotype; one-way ANOVA followed by Tukey’s post hoc test; NS, not significant). Scale bar, 50 μm. (**D**) A Venn diagram showing the 31 overlapping proteins between the 132 RBPs listed in the RBPmap database and the 1,334 DEPs identified in *Cyfip2^+/R87C^* mice. The graph below shows the *P*-values and fold changes of the 31 RBPs; FMRP and PUM1 are highlighted in bold. Right panels show representative Western blot images and their quantification for CYFIP2, WAVE1, PUM1, and FMRP in the hippocampal CA1 region of WT and *Cyfip2^+/R87C^*mice at PWK 7 (n = 8 animals per genotype; unpaired two-tailed Welch’s t-tests; numbers in parentheses indicate the molecular weight [kDa] of each protein, and GAPDH was used as a loading control). Data are presented as mean ± SEM. \**P* < 0.05, \*\**P* < 0.01, \*\*\**P* < 0.001.

We reasoned that proteins whose abundance is altered as a consequence of RBP-mediated changes in translation would be captured in our quantitative global proteomic dataset (Fig. 4B). To identify such candidates, we first selected DEPs from the 1,334 DEPs identified in the hippocampal CA1 region of *Cyfip2^+/R87C^* mice using the criteria of *P* < 0.01 and |log2 fold change| > 0.2, and then excluded WRC components and RBPs, which are considered to be directly affected by CYFIP2 Arg87Cys (Fig. 6B, left panel). This filtering strategy yielded 12 up-regulated and 21 down-regulated DEPs. Among these candidates, Eph receptor A3 (EphA3) emerged as a particularly interesting protein with respect to cell adhesion and migration (Fig. 6B, right panel). EphA3 is a member of the receptor tyrosine kinase family that binds ephrin ligands and mediates bidirectional signaling, thereby regulating processes such as cell-cell adhesion and cell motility (Pasquale, 2008; Singh et al., 2012). Notably, in the hippocampal CA1 region, *EphA3* mRNA is preferentially expressed in superficial-layer pyramidal neurons (Supplementary Material, Fig. S4), suggesting a potential role in modulating adhesion or migration-related properties of this neuronal population.

To confirm the reduction of EphA3 expression in *Cyfip2^+/R87C^* hippocampal CA1 pyramidal neurons, we performed immunofluorescence staining for EphA3. Because EphA3 mRNA is preferentially expressed in superficial-layer pyramidal neurons, we co-stained sections with CalB to identify this neuronal population. In WT mice, nearly all CalB-positive neurons (∼100%) were located within 50 μm of the boundary between the stratum pyramidale and stratum radiatum, consistent with the typical distribution of superficial CA1 pyramidal neurons. In contrast, in *Cyfip2^+/R87C^* mice, approximately 80% of CalB-positive neurons were located within this 50 μm zone, whereas the remaining ∼20% were distributed beyond this distance, reflecting the neuronal layer dispersion observed in these mice. Based on this distribution pattern, we compared EphA3 immunofluorescence intensity across three groups of neurons: (i) CalB-positive neurons within 50 μm of the boundary in WT mice, (ii) CalB-positive neurons located within 50 μm of the boundary in *Cyfip2^+/R87C^* mice, and (iii) CalB-positive neurons located >50 μm from the boundary in *Cyfip2^+/R87C^* mice. Quantitative analysis revealed that EphA3 intensity was significantly reduced in both *Cyfip2^+/R87C^* neuronal groups compared with WT neurons (Fig. 6C). Within the *Cyfip2^+/R87C^* mice, neurons located >50 μm from the boundary tended to show a greater reduction in EphA3 intensity than those within 50 μm; however, this difference did not reach statistical significance.

Next, we sought to identify candidate RBPs that may be involved in the translational repression of *EphA3* mRNA in *Cyfip2^+/R87C^* hippocampal CA1 pyramidal neurons. To this end, we utilized RBPmap (Paz et al., 2014) to predict RBPs with potential binding sites on mouse *EphA3* mRNA. We first compared the 132 RBPs available in the RBPmap database with our DEP list from the *Cyfip2^+/R87C^* hippocampal CA1 region, identifying 31 overlapping RBPs (Fig. 6D). As expected, all 31 RBPs were up-regulated DEPs. Among them, U2 small nuclear RNA auxiliary factor 2 (U2AF2), fragile X messenger ribonucleoprotein (FMRP), PUM1, and poly(rC)-binding protein 3 (PCBP3) were particularly prominent based on their statistical significance and fold change. Among these candidates, FMRP and PUM1 are well-known translational repressors of target mRNAs (Darnell et al., 2011; Gennarino et al., 2015; Van Etten et al., 2012), making them especially relevant to our hypothesis. Immunoblot analysis of the hippocampal CA1 region confirmed the reduced levels of CYFIP2 and WAVE1 detected in our proteomic analysis, and further confirmed that PUM1 protein levels were significantly increased in *Cyfip2^+/R87C^*mice compared to WT controls (Fig. 6D). Although FMRP also showed an increasing trend, this change did not reach statistical significance. In addition, RBPmap-based prediction indicated that PUM1 has a high likelihood (*P* < 0.0001) of binding to *EphA3* mRNA (Supplementary Material, Table S12). Taken together, these findings raise the possibility that increased PUM1 expression in *Cyfip2^+/R87C^* hippocampal CA1 pyramidal neurons may contribute to reduced EphA3 protein levels through translational repression of its mRNA. However, this interpretation remains speculative, and direct experimental validation will be required to establish a causal relationship.

## Discussion

In this study, we sought to elucidate the molecular mechanisms underlying the toxic gain-of-function effects of the *CYFIP2* Arg87Cys variant. Given that CYFIP2 is predominantly expressed in neurons rather than glial cells (Ma et al., 2022; Y. H. Zhang et al., 2019), we focused on the layer dispersion of CA1 pyramidal neurons observed at PWK 7, a developmental stage preceding overt gliosis in the hippocampus of *Cyfip2^+/R87C^* mice. Exploratory snRNA-seq of the hippocampus generated cell type-specific hypotheses implicating altered cell adhesion- and migration-related pathways in CA1 pyramidal neurons, consistent with the observed structural phenotype. In parallel, HEK293T cells harboring the *CYFIP2* Arg87Cys variant exhibited reduced proliferation and morphological abnormalities, accompanied by concordant changes in the expression of cell adhesion-related genes as assessed by RNA sequencing. Importantly, quantitative global proteomic analysis of the CA1 region from *Cyfip2^+/R87C^* mice identified a marked increase in RBPs alongside a reduction in components of the WRC, suggesting that an imbalance between these two functional protein groups may represent a key feature of the CYFIP2 Arg87Cys-associated phenotype. Additional phosphoproteomic analysis further indicated potential functional alterations in synaptic proteins, pointing to changes in synaptic regulation, and electrophysiological and morphological analyses of CA1 pyramidal neurons revealed increased mEPSC frequency and altered dendritic architecture, most prominently in the more severely dispersed deep-layer population. Finally, increased PUM1 expression may be associated with reduced EphA3 protein levels, potentially through effects on mRNA translation, and could be related to CA1 pyramidal neuron dispersion. Collectively, our findings suggest that the *CYFIP2* Arg87Cys variant perturbs neuronal architecture and function through coordinated dysregulation of cytoskeletal regulation, mRNA processing/translation, and synaptic signaling pathways, in a manner that is distinct from the consequences of *CYFIP2* loss-of-function (Ma, Kim, et al., 2025; Y. Zhang et al., 2019).

How expression of the CYFIP2 Arg87Cys protein leads to a reduction in WRC components while simultaneously increasing RBPs remains enigmatic, particularly in light of our previous finding that CYFIP2 Arg87Cys protein undergoes hyperubiquitination and proteasomal degradation (Kang et al., 2023; Lee et al., 2019). Structural predictions based on available WRC crystal structures indicate that the Arg87 residue of CYFIP2 lies within the interaction interface between CYFIP2 and WAVE proteins (Chen et al., 2010; Nakashima et al., 2018). Accordingly, the Arg87Cys substitution is predicted to weaken CYFIP2-WAVE interactions, potentially promoting dissociation of CYFIP2 from the WRC and subsequent destabilization of WRC components including CYFIP2 itself. However, it is unlikely that all WRC-dissociated CYFIP2 Arg87Cys protein is immediately degraded; instead, a fraction may escape proteasomal degradation and become more available to interact with alternative binding partners, such as RBPs, which associate with CYFIP2 independently of the WRC. One speculative possibility is that this residual CYFIP2 Arg87Cys preferentially associates with RBPs. Such interactions could stabilize the remaining CYFIP2 Arg87Cys protein while concurrently enhancing the stability of its RBP partners.

Given that many RBPs form extensive physical and functional interaction networks within the cell (Hentze et al., 2025; Street et al., 2024), CYFIP2 Arg87Cys may not need to directly interact with all RBPs found to be increased in our global proteomic analysis. Rather, interactions with a limited set of core RBPs could influence broader RBP complexes, thereby secondarily increasing the abundance of multiple RBPs. In this context, RBPs such as Ago and PUM1, previously identified in the CYFIP2 brain interactome (Y. Lee et al., 2020), may represent candidate primary interactors. In addition, some of the increased RBPs may reflect secondary effects, for example through enhanced translation of their mRNAs driven by initially up-regulated RBPs (Khoroshkin et al., 2024). Beyond effects on protein abundance, association of the CYFIP2 Arg87Cys protein with RBPs may also influence their intracellular localization, interactions with target mRNAs, or binding affinities, thereby further modulating RBP function (Kedersha et al., 2016; Pekovic et al., 2025; Smith et al., 2021). Although these possibilities are not readily testable at present, future studies aimed at defining the interactome specific to the CYFIP2 Arg87Cys protein, together with approaches such as ribosome profiling in *Cyfip2^+/R87C^* mice, should help elucidate the mechanisms underlying RBP up-regulation and its consequences for mRNA processing and translation in these mice.

Phosphoproteomic analysis, together with GO enrichment of proteins exhibiting altered phosphorylation, suggested functional alterations in glutamatergic synaptic proteins in CA1 pyramidal neurons of *Cyfip2^+/R87C^* mice. Notably, among the top-ranked phosphoproteins, Syn1 and Bsn play essential roles in presynaptic active zone organization and neurotransmitter release (Sudhof, 2012). In addition, SynGAP1 is known to interact with multiple scaffolding and signaling proteins at the postsynapse and to critically regulate both structural and functional synaptic plasticity (Gamache et al., 2020). These observations raise the possibility that altered phosphorylation of synaptic proteins may contribute to changes in synaptic transmission and plasticity in CA1 pyramidal neurons of *Cyfip2^+/R87C^* mice. Consistent with this possibility, our whole-cell recordings revealed an increased frequency of mEPSCs without a corresponding change in mEPSC amplitude, a pattern classically associated with presynaptic rather than postsynaptic alterations. Because we detected no differences in vGluT1/Homer1 puncta density or in the abundance of representative synaptic proteins at PWK 7, the increased mEPSC frequency is unlikely to reflect a simple increase in the number of excitatory synapses, and may instead arise from altered presynaptic release properties, in line with the altered phosphorylation of Syn1 and Bsn.

Beyond synaptic regulation, it is noteworthy that several highly ranked phosphoproteins are also associated with mRNA translation and protein synthesis. This raises the intriguing possibility that phosphorylation changes in these proteins may be functionally linked to the RBP up-regulation observed in *Cyfip2^+/R87C^* mice. For example, Rplp2 is a core component of the ribosome, while SynGAP1 negatively regulates protein synthesis through modulation of the Ras-extracellular signal-regulated kinase (ERK) and mechanistic target of rapamycin complex 1 (mTORC1) signaling pathways (Wang et al., 2013). Map2 has been reported to interact with multiple ribosomal components in the mouse brain (Grubisha et al., 2021), and Mark4 functions as a negative regulator of mTORC1 activity (Li & Guan, 2013; Samra et al., 2024). In addition, Dgkβ has been implicated in mTORC1 regulation through the generation of phosphatidic acid (Nakai et al., 2020). Taken together, given the intimate coupling between synaptic plasticity and protein synthesis (Klann & Dever, 2004), these findings underscore the need for an integrated understanding of how phosphorylation-dependent regulation of synaptic and translational machinery converges to shape neuronal function in *Cyfip2^+/R87C^* mice.

Our morphological analysis of the recorded neurons revealed that dendritic architecture was also altered, with reduced total apical dendritic length in deep-layer neurons and a redistribution of basal dendritic branching, characterized by increased proximal and decreased distal intersections, in both sublayers. Notably, these structural changes were accompanied by an increase, rather than a decrease, in mEPSC frequency, further arguing against a simple relationship between dendritic size and the number of functional excitatory synapses. Instead, the proximal shift in basal dendritic branching could alter the spatial distribution of excitatory inputs along the dendritic arbor, which may in turn influence synaptic integration and, together with altered presynaptic properties, contribute to the observed changes in mEPSC frequency.

A notable feature of these synaptic and dendritic phenotypes is their laminar specificity: both the increase in mEPSC frequency and the dendritic alterations were most pronounced in deep-layer neurons, which also exhibit more severe dispersion than superficial-layer neurons. Superficial and deep CA1 pyramidal neurons are known to differ in their gene expression profiles, afferent connectivity, and intrinsic and synaptic properties (Soltesz & Losonczy, 2018), and the laminar position of a pyramidal neuron is a key determinant of the inputs it receives. We therefore speculate that the more severe mispositioning of deep-layer neurons in *Cyfip2^+/R87C^* mice may secondarily perturb their dendritic organization and excitatory input, thereby amplifying the synaptic and dendritic phenotypes in this population. At the same time, it is important to note that these phenotypes cannot be attributed to altered positioning alone. The WRC is a central regulator of actin dynamics in dendrites and spines (Choi & Han, 2015; Ethell & Pasquale, 2005; Soderling et al., 2007), and the reduction of WRC components that we observed in *Cyfip2^+/R87C^* mice would be expected to directly affect dendritic growth, branching, and synapse formation, independently of neuronal positioning. Distinguishing the relative contributions of altered positioning, WRC reduction, and RBP-mediated translational dysregulation to these synaptic and dendritic phenotypes will require approaches that manipulate these processes separately, such as cell type- or sublayer-specific rescue experiments.

The hippocampal CA1 pyramidal neuron layer is established through highly coordinated molecular mechanisms that regulate neuronal migration, positioning, and subsequent laminar organization. Classical pathways implicated in this process include Reelin-Disabled-1 (Dab1) signaling, in which Dab1 functions as a key intracellular adaptor downstream of Reelin receptors, as well as microtubule- and dynein-dependent migration machinery involving platelet-activating factor acetylhydrolase 1b regulatory subunit 1 (Pafah1b1; also known as Lissencephaly 1 [Lis1]), nuclear distribution element-like 1 (Ndel1), doublecortin (Dcx), and related cytoskeletal regulators (Alexander et al., 2023; Corbo et al., 2002; Garrott et al., 2022; Reiner & Sapir, 2013). Disruption of these pathways has been shown to result in abnormal lamination and dispersion of pyramidal neurons within the hippocampus. Importantly, accumulating evidence indicates that these molecular mechanisms are not restricted to embryonic development but continue to play essential roles in maintaining hippocampal layer integrity during postnatal life and adulthood. Notably, postnatal conditional deletion of *Ndel1* using calcium/calmodulin-dependent protein kinase II alpha (*CaMKII*α)-Cre results in age-progressive dispersion of CA1 pyramidal neurons, indicating an ongoing requirement for cytoskeletal and Reelin-associated signaling to maintain CA1 layer compaction after initial lamination (Jiang et al., 2016). Interestingly, these two classes of mechanisms may be mechanistically linked at the level of CYFIP proteins. Ndel1 directly binds CYFIP1 and HSPC300, and is required for WRC assembly by stabilizing nascent subcomplexes: its depletion reduces the level of mature WRC and abolishes WRC-dependent actin polymerization (Wu et al., 2012). Although it remains unknown whether Ndel1 similarly engages CYFIP2, this raises the possibility that the reduced WRC levels in *Cyfip2^+/R87C^* mice and the Ndel1-dependent maintenance of CA1 layer compaction converge on a shared actin-regulatory process. In this context, the progressive postnatal dispersion of CA1 pyramidal neurons in *Cyfip2^+/R87C^*mice is unlikely to arise from a single molecular alteration, but may instead reflect the convergence of multiple disrupted pathways involved in layer maintenance. While we propose reduced EphA3 expression as one potential contributor to impaired cell-cell adhesion or neuronal positioning, we cannot exclude the possibility that perturbations in these well-established layer maintenance pathways act in parallel or synergistically to drive the observed phenotype. In addition, the reduction of WRC components in *Cyfip2^+/R87C^* mice is expected to alter actin cytoskeletal dynamics, which are critical for maintaining neuronal morphology and intercellular packing within the pyramidal layer. Together, these considerations underscore the need for an integrated mechanistic framework in which EphA3 signaling, Reelin- and Ndel1-associated cytoskeletal regulation, and WRC-dependent actin remodeling collectively contribute to the progressive destabilization of the CA1 pyramidal neuron layer.

A further consideration concerns the sublayer specificity of the candidate mechanisms we propose. *Epha3* mRNA is expressed almost exclusively in superficial-layer CA1 pyramidal neurons (Supplementary Material, Fig. S4), and accordingly we quantified EphA3 immunoreactivity in CalB-positive neurons, which correspond to this population. Reduced EphA3 levels are therefore unlikely to act cell-autonomously in deep-layer neurons. However, our quantification of cell distribution along the radial axis indicated that dispersion is more pronounced in CalB-negative, deep-layer neurons, and the synaptic and dendritic alterations described above were likewise most prominent in this population. Two non-exclusive explanations may reconcile this apparent discrepancy. First, because Eph receptors and ephrins mediate bidirectional signaling between neighboring cells (Pasquale, 2008), reduced EphA3 in superficial-layer neurons could non-cell-autonomously affect the adhesive interactions and positioning of adjacent deep-layer neurons. Second, additional molecular mechanisms may operate within deep-layer neurons themselves. In this regard, it is notable that the up-regulation of PUM1 and other RBPs was detected in CA1 tissue as a whole rather than in a sublayer-restricted manner, raising the possibility that other PUM1 target mRNAs, or targets of other up-regulated RBPs, mediate comparable effects on adhesion- and cytoskeleton-related proteins in deep-layer neurons. Distinguishing between these possibilities will require sublayer-resolved approaches, such as spatial transcriptomic or proteomic profiling combined with sublayer-specific manipulation of candidate molecules.

Notably, in *Cyfip2^+/R87C^* mice, neuronal layer dispersion precedes the emergence of pronounced neuronal loss and gliosis, which become evident at later stages and coincide temporally with the onset of spontaneous recurrent seizures after PWK 14. This temporal sequence raises the possibility that early structural disorganization of the CA1 pyramidal layer, together with the accompanying synaptic and dendritic alterations described above, constitutes a permissive or initiating event that predisposes the hippocampal network to subsequent neurodegeneration, glial activation, and epileptogenesis (Ma, Kang, et al., 2025). Defining the precise molecular mechanisms underlying this early dispersion phenotype and determining whether its prevention can attenuate later cellular pathology and seizure development, therefore represents a critical question for future studies.

In conclusion, these findings support a toxic gain-of-function model for the *CYFIP2* Arg87Cys variant and highlight the importance of variant-specific molecular characterization in understanding *CYFIP2*-associated disease, underscoring the need for integrated, multi-level experimental approaches to dissect pathogenic mechanisms of disease-associated missense variants.

## Materials and Methods

### Animals

*Cyfip2^+/R87C^* and *Cyfip2^+/-^* mice (C57BL/6N background) used in this study have been described previously (Kang et al., 2023; S. H. Lee et al., 2020; Zhang et al., 2018). Only male mice were used to maintain consistency across experimental cohorts and with our previous characterization of this mouse model, thereby reducing heterogeneity in this mechanistic study. In total, 98 mice were used for all experiments. Unless otherwise indicated, all experiments were performed using 7-week-old (PWK 7) mice. For the experiment shown in Figure 1A, 1-week-old (PWK 1) mice were used. Body weight at the time of experimentation was 23.5 ± 1.8 g for WT mice, 21.6 ± 2.6 g for *Cyfip2^+/R87C^* mice, and 24.3 ± 1.4 g for *Cyfip2^+/-^* mice at PWK 7 (mean ± SD). Mice were group-housed (4–5 animals per cage) in standard cages under controlled temperature and humidity with a 12-h light/12-h dark cycle and had ad libitum access to standard chow and water. Mice were anesthetized with inhaled isoflurane in a closed induction chamber prior to terminal tissue collection. Isoflurane was applied to an absorbent material within the chamber to allow vapor inhalation, and adequate anesthesia was confirmed by loss of postural reflex before tissue harvesting. Animals were then euthanized by cervical dislocation as a secondary physical method, followed by rapid decapitation for brain tissue collection.

### Fluorescence immunohistochemistry

Fluorescence immunohistochemistry and analyses were conducted as described previously (B. Lee et al., 2017; Ma, Kang, et al., 2025). Mice were anesthetized with isoflurane and transcardially perfused with heparinized (20 units/ml) phosphate-buffered saline (PBS), followed by 4% paraformaldehyde (PFA) in PBS. Brains were then extracted and post-fixed overnight in 4% PFA. After post-fixation, brain tissue was washed with PBS and cryoprotected in 30% sucrose in PBS for 48 h. The brain tissues were frozen in O.C.T compound (SAKURA Tissue-Tek, catalogue number 4583) and coronally sectioned at 60 μm using a cryostat microtome (Leica, CM3050S). Sections were collected from the following approximate Bregma coordinates: hippocampus: −1.555 mm to −2.355 mm. The samples were washed with 0.1% Triton X-100 in PBS and blocked with PBS containing 3% bovine serum albumin and 0.5% Triton X-100. Confocal microscopy (Zeiss, LSM900) was used for image acquisition. Z-stack images were acquired as follows: 5 optical sections spanning a total thickness of 40 μm for parvalbumin, 15 optical sections spanning 7 μm for vGluT1 and Homer1, and 9 optical sections spanning 8 μm for EphA3 and Calbindin. For analysis, 3D reconstructions were generated using Imaris 10.0.1 (Bitplane, UK), and the number and intensity of puncta were measured by spot function. Somatosensory cortical regions were imaged by tile-scanning confocal microscopy, and whole hippocampal regions were acquired using a slide scanner (Zeiss Axio Scan.Z1). The primary antibodies used are listed in Table S13.

### Western blot analysis

Brain lysate preparation and Western blot analyses were conducted as described previously (Jin et al., 2019; Jin et al., 2021). Briefly, mouse brain samples were homogenized in RIPA buffer (50 mM Tris–HCl pH 8.0, 150 mM NaCl, 0.1% SDS, 1% Triton X-100, and 0.5% sodium deoxycholate) with freshly added protease and phosphatase inhibitors (Sigma-Aldrich, catalogue number 05892970001 and 04906837001, respectively). The homogenate was centrifuged at 13,000 rpm for 15 min at 4°C, and the resulting supernatant was transferred to new tubes for further processing in Western blotting. Protein concentrations were determined using the Bio-Rad Protein Assay (Bio-Rad, catalogue number 5000006). From each sample, 10 μg of protein was loaded onto 4-15% Mini-PROTEAN TGX Precast Protein Gels (Bio-Rad, catalogue number 4561084). The proteins were then transferred to a nitrocellulose membrane (GE Healthcare, catalogue number 10600001). Tris-buffered saline with Tween 20 (TBST, 0.05%) solution was used for washing, and 5% skim milk in TBST was used for blocking. The Western blot images were captured using the ChemiDoc Touch Imaging System (Bio-Rad) and analyzed for quantification using ImageJ software. The primary antibodies used are listed in Table S13.

### Gene editing in HEK293T cells

To prepare the CRISPR/Cas9 system, the single-guide RNA (sgRNA) was designed using CRISPR RGEN Tools (http://www.rgenome.net/). The sgRNA construct was cloned into the pGL3-U6-sgRNA-EGFP vector at the BsaI restriction enzyme site. For Cas9 expression, the p3S-spCas9 plasmid was utilized. Single-stranded oligo deoxynucleotide (ssODN) was synthesized by Integrated DNA Technologies (IDT) as a donor template for homology-directed repair (HDR), including the desired genetic modification. HEK293T cells were cultured in Dulbecco’s Modified Eagle Medium (DMEM) supplemented with 10% fetal bovine serum (FBS) and maintained in a CO_2_ incubator at 37 °C. Lipofectamine 3000 (Invitrogen, catalogue number L3000015) was used for transfection. One day prior to transfection, 2 × 10^4^ cells were seeded into a 24-well plate. For transfection, 750 ng of sgRNA, 250 ng of spCas9, and 1000 ng of donor template were used, following the manufacturer’s protocol. After 72 h, the cells were harvested, and green fluorescent protein (GFP)-positive cells were sorted into a 96-well plate using fluorescence-activated cell sorting (FACS). The sequences of sgRNA and ssODN utilized are listed in Supplementary Material, Table S14. After sufficient cell expansion, genomic DNA was extracted from the cells in each well. The genomic DNA was amplified for the target region using nested PCR for genotyping and targeted deep sequencing analysis. The amplified DNA was then analyzed for nucleotide sequences using the MiSeq sequencing system (Illumina). Illumina sequencing data were subsequently processed using the EUN program (https://daeunyoon.com/). All primers utilized are listed in Supplementary Material, Table S15. Off-target sites were identified in exons of the Homo sapiens (GRCh38/hg38) genome using Cas OFFinder from CRISPR RGEN Tools (http://www.rgenome.net/), permitting up to three mismatches. Off-target events were assessed by amplifying genomic DNA, followed by targeted deep sequencing. The *CYFIP2* off-target primer sets utilized are listed in Supplementary Material, Table S15.

### Cell proliferation and morphology analysis

HEK293T cells were monitored for 96 h post-seeding using the EVOS M5000 imaging system (Thermo Fisher Scientific). Cellular confluency (%) was automatically quantified using the confluency tool within the EVOS M5000. Cell roundness and area were analyzed from the acquired images using ImageJ software, with cell boundaries defined by manual outlining of individual cells. Experiments were independently repeated three times using separate cell culture preparations from different passages. For each independent experiment, confluency was obtained from a single field per genotype, and roundness and area were averaged across the cells measured in that experiment; the mean value of each independent experiment was used as a single data point for statistical analysis.

### 3D Holotomography imaging, data acquisition, and quantitative analysis

3D holotomography was performed using an HT-X1 system (Tomocube Inc., Republic of Korea) employing low-coherence illumination to minimize coherent noise. A 450 nm light-emitting diode source and a 40× motorized objective (NA 0.95) were used, with illumination patterns modulated by a digital micromirror device and delivered through a condenser lens (NA = 0.72, working distance = 30 mm). Imaging was performed under controlled conditions at 37°C and 5% CO using a stage-top incubator (Tokai Hit, Japan). For 3D refractive index (RI) tomogram acquisition, cells were captured across multiple axial planes within a 165 μm × 165 μm field of view using TomoStudio X software (Tomocube Inc.). Focus was maintained via a 780 nm laser-based autofocus system. RI tomograms were reconstructed with deconvolution algorithms, yielding lateral and axial resolutions of 155 nm and 947 nm, respectively. Cell 3D morphology was visualized in TA Viewer (Tomocube Inc.), and cellular sphericity and volume were quantified in TomoAnalysis (Tomocube Inc.) after RI-based cell segmentation with morphological refinement. Imaging was performed in three independent experiments using separate cell culture preparations from different passages, and 21–33 cells per genotype were analyzed in total. Each cell was treated as an individual data point for statistical analysis.

### Bulk RNA sequencing and analysis of HEK293T cells

Bulk RNA sequencing and downstream analyses of HEK293T cells were performed as described previously (Ma, Kang, et al., 2025). Total RNA extraction, library preparation, cluster generation, and sequencing were conducted by Macrogen Inc. (Seoul, Korea). RNA-seq libraries were prepared using the TruSeq Stranded Total RNA LT Sample Prep Kit (Illumina) according to the manufacturer’s instructions. Sequencing was performed on an Illumina platform to generate 101-bp paired-end reads.

Transcript abundance was estimated using the pseudo-alignment–based method implemented in Salmon (v1.1.0) (Patro et al., 2017), with reads mapped to the human reference genome (GRCh38) using GENCODE v32 gene annotations. Differential gene expression analysis was performed using DESeq2 (v1.44.0) (Love et al., 2014). *P* values were adjusted for multiple testing using the Benjamini–Hochberg method, and genes with an adjusted *P* value < 0.05 and an absolute fold change greater than 1.5 were considered differentially expressed.

Gene Ontology (GO) and Kyoto Encyclopedia of Genes and Genomes (KEGG) pathway enrichment analyses were performed using DAVID (v2021; https://davidbioinformatics.nih.gov/) (Dennis et al., 2003), with down-regulated and up-regulated DEGs analyzed separately against a background of all genes detected in the RNA-seq dataset. For each category, the five top-ranked terms are shown in Figure 3E, and terms with a Benjamini-Hochberg-adjusted *P* value < 0.05 were considered significantly enriched. The raw data have been submitted to the Gene Expression Omnibus (GEO) repository under the accession number GSE317553.

### Single-nucleus RNA sequencing data generation and analysis

Nuclei were isolated from hippocampal tissue samples using the Singulator™ 100 system (S2 Genomics), which automatically processes tissues into suspensions of intact nuclei. Cellular debris was removed using a Percoll density gradient, and nuclei concentration was assessed with the LUNA-FL™ Automated Fluorescence Cell Counter (Logos Biosystems) following the recommendations provided in the 10× Genomics cell preparation guidelines. Single-nucleus RNA-seq libraries were prepared using the Chromium X platform (10× Genomics) according to the Chromium GEM-X Single Cell 3′ Reagent Kit v4 protocol (CG000731). Libraries were quantified by qPCR (KAPA) and assessed using the Agilent 4200 TapeStation, then sequenced on an Illumina NovaSeq 6000 platform according to the recommended read configuration.

Raw sequencing data were processed using the 10× Genomics Cell Ranger pipeline (v 8.0.1) to generate gene-barcode count matrices. Downstream analyses were performed in R using the Seurat package (v5.1.0) (Stuart et al., 2019). Low-quality nuclei and technical artifacts were removed through standard quality-control procedures, including filtering based on transcript complexity and mitochondrial gene content, and putative doublets were identified and excluded using DoubletFinder (v 2.0.6) (McGinnis et al., 2019). Specifically, nuclei with fewer than 500 or more than 6,000 detected genes were excluded, and only nuclei with no detectable mitochondrial gene expression were retained to ensure high-quality nuclear transcriptomes, as mitochondrial RNA is largely absent in intact nuclei from hippocampal tissue and may indicate cytoplasmic contamination or damaged cells. Rarely expressed genes were filtered out prior to downstream analyses, and the remaining nuclei and genes were retained for further processing.

Filtered datasets were integrated across samples, and batch effects were corrected by applying Harmony (v1.2.3) (Korsunsky et al., 2019). The resulting low-dimensional representation was used for clustering and visualization with uniform manifold approximation and projection (UMAP) in Seurat (Stuart et al., 2019). Cluster-specific marker genes were identified using differential expression analysis based on the MAST statistical framework (Finak et al., 2015). Cell clusters were annotated based on the expression patterns of representative marker genes observed in the present dataset, in combination with reference-based annotation using SingleR (v2.4.1) (Aran et al., 2019). Major neuronal and glial cell types were identified in a manner consistent with previous single-nucleus RNA sequencing studies of the hippocampus (Cid et al., 2021; Takahashi et al., 2024; Zhu et al., 2024), resulting in the identification of eleven distinct cell types. Excitatory neuronal subtypes corresponding to hippocampal CA1-3 regions and the dentate gyrus were distinguished. Exploratory differential expression analyses between experimental groups were performed using MAST, both across all nuclei and within annotated cell types, to generate candidate transcriptional signatures for downstream pathway analysis. Genes with a false discovery rate (FDR) < 0.05 were considered candidate differentially expressed genes (DEGs). Functional interpretation of candidate DEGs was performed by over-representation analyses including Enrichr (v3.4) (Chen et al., 2013) with the Gene Ontology Biological Process, Molecular Function, and Cellular Component 2023 libraries. For each cell type, candidate up-regulated and down-regulated DEGs were analyzed separately, and *P* values were adjusted using the Benjamini-Hochberg method. Gene sets with adjusted *P* < 0.05 are listed in Supplementary Material, Table S2. For Excitatory-CA1 neurons, the five top-ranked terms in each Gene Ontology category are shown in Figure 2C, irrespective of statistical significance, for exploratory purposes. The raw data have been deposited in the GEO under the accession number GSE317598.

### Sample preparation for proteomic and phosphoproteomic analysis

Hippocampal CA1 regions were isolated from WT and *Cyfip2^+/R87C^* brains at PWK 7 and homogenized and lysed with the 1× sodium dodecyl sulphate (SDS) buffer (5% SDS, 50 mM triethylamonium bicarbonate (TEAB), pH 8.5). 300 μg of lysates were purified and digested with S-trap mini digestion kit (ProtiFi, Huntington, NY) following the provided protocol. In this case, 10 μg amounts of mass-spectrometry-grade Trypsin Gold (Promega, Madison, WI) were used for each sample (1:30). Tryptic digested peptide were dried and quantified with a Pierce quantitative colorimetric peptide assay kit (Thermo Fisher Scientific, Rockford, IL). 100 μg of trypsin-digested peptides from each sample were labeled with 18-plex tandem mass tag (TMT) isotopes reagents (Thermo Fisher Scientific, Rockford, IL) according to the manufacturer’s protocol. Peptides from WT mice were labeled with the 126, 127N, 127C, 128N, 128C, 129N, 129C, 130N tags and peptides from *Cyfip2^+/R87C^* mice were labeled with the 130C, 131N, 131C, 132N, 132C, 133N, 133C, 134N tags. For quality control in TMT experiments, reference samples were prepared by pooling of same amount of all samples and labeled with 134C and 135N TMT tags. Each TMT channel was freshly dissolved in anhydrous acetonitrile (ACN) with a ratio of 0.5:20 (w:v, mg:mL). After incubation for 1 h at room temperature, the reaction was quenched by adding 5 μL of 5% hydroxylamine to each sample, followed by incubation for 15 min. Then, samples were combined, dried, and desalted with Pierce peptide desalting spin columns (Thermo Fisher Scientific, Rockford, IL). Total peptides were fractionated into 20 fractions by basic reverse phase liquid chromatography and each eluted peptide sample was vacuum dried. For parallel analyses, 5% of the TMT-labeled peptide mixture was used for total proteome profiling, whereas the remaining 95% was reserved for phosphoproteomic analysis.

The phosphoproteomic fractions were concatenated from 20 into 10 fractions for phosphopeptide enrichment experiments. For phosphopeptide enrichment, 30 μL of PTMScan IMAC Fe-NTA magnetic beads were washed three times with deionized water and then incubated overnight at 4°C on a rotator with each of the 10 fractionated samples in 80% ACN in 0.1% trifluoroacetic acid (TFA). After overnight incubation, the beads were washed three times with 80% ACN in 0.1% TFA, and the phosphopeptides attached to the beads were eluted using an elution buffer (50% ACN in 1% ammonium hydroxide). Following elution, the phosphopeptides were promptly acidified to a pH of 3.5–4.0 using 10% TFA before undergoing vacuum drying. The samples were subsequently desalted using a C18 disk, followed by gradient elution with 40% acetonitrile containing 0.1% formic acid. All eluates were then dried using a SpeedVac.

#### LC-MS/MS and protein identification for proteomic analysis

LC-MS/MS analysis was performed using an UltiMate 3000 RSLCnano system (Thermo Scientific) coupled to an Orbitrap Exploris 480 mass spectrometer (Thermo Fisher Scientific) at the Korea Basic Science Institute (KBSI). Samples were trapped on 75 μm × 2 cm C18 precolumn (nanoViper, Acclaim PepMap100, Thermo Fisher Scientific) before being separated on an analytical C18 column (75 μm × 50 cm PepMap RSLC, Thermo Fisher Scientific) for 140 min at a flow rate of 250 nL/min. The mobile phase A consisted of 0.1% formic acid in water, and mobile phase B consisted of 99.9% acetonitrile containing 0.1% formic acid. The LC gradient started with 5% B for 20 min. It was ramped to 13% B buffer for 50 min followed by a gradient from 13 to 25% B buffer for 65 min, then 25 to 95% for 5 min, and 95% for 5 min, followed by column equilibration at 5% B buffer for 15 min prior to the next run. During the chromatographic separation, the Orbitrap Exploris 480 was operated in data-dependent mode, automatically switching between MS1 and MS2. The full scan resolution was 120 000 at m/z 400. MS2 scans were performed using higher-energy collisional dissociation (HCD) with normalized collision energies of 37.5% for global proteomics and 30.0% for phosphoproteomics. Previously fragmented ions were excluded for 30 s within 10 ppm. The electrospray voltage was maintained at 1800 V, and the capillary temperature was set to 275°C.

MS/MS spectra were identified and quantified using Integrated Proteomics Pipeline software (Integrated Proteomics Applications, Inc., San Diego, CA, USA) with the Uniprot mouse database (Release 2025). Search parameters included a precursor mass tolerance of 10 ppm, a fragment ion mass tolerance of 50 ppm, and a minimum of two peptides required for protein identification at a false discovery rate less than 0.01. TMT reporter ion quantification was performed with Census (Park et al., 2014) implemented in the Integrated Proteomics Pipeline software with a mass error tolerance of 20 ppm. Statistical analysis was performed with Perseus (Tyanova et al., 2016) software (version 1.6.15). Protein expression levels between samples were compared using Welch’s t-test, with *P* value < 0.05 considered statistically significant. All raw MS data files from this study have been deposited to the MassIVE repository with the ProteomeXchange identifier PXD073375.

### Bioinformatic analysis of proteomic and phosphoproteomic data

Proteins with a Welch’s t-test *P* value < 0.05 were defined as DEPs, as described above. Enriched terms among the 1,334 DEPs and the relationships between them were analyzed using Metascape (https://metascape.org), and the results were visualized as an enrichment network in which each node represents an enriched term and nodes belonging to the same cluster share a color. GSEA was conducted using the GSEAPreranked module (gsea-4.3.3.jar) (Subramanian et al., 2005) with gene sets obtained from the Molecular Signatures Database (MSigDB, v2023.2.Hs). Up-regulated and down-regulated proteins were analyzed separately, irrespective of statistical significance, and the results are presented as normalized enrichment scores (NES). GO and KEGG pathway enrichment analyses were performed separately for the 693 up-regulated and 641 down-regulated DEPs, and the same approach was applied to the phosphoproteins showing increased or decreased phosphorylation. *P* values were adjusted for multiple testing using the Benjamini–Hochberg method. For each GO category and for KEGG, the five top-ranked terms are presented; terms with an adjusted *P* value < 0.05 are indicated by solid bars, and terms not reaching this threshold by light-colored bars. For the comparison with the previously reported *Cyfip2* KO proteomic dataset (Zhang et al., 2024), over-representation analysis of GO terms was performed separately for the 18 DEPs that were up-regulated in *Cyfip2*^+/R87C^ mice but down-regulated in *Cyfip2* KO mice and for the 19 DEPs that were down-regulated in both models. Because the two groups yielded different numbers of enriched terms, an FDR < 0.05 was applied to the former and a more stringent threshold (FDR < 0.02, with at least three assigned proteins per term) to the latter.

### Phosphopeptide-global protein log fold change comparative analysis

Changes in phosphopeptide abundance were evaluated relative to their corresponding protein levels by comparing phosphopeptide log fold change with global protein log fold change. Scatter plots were generated with Phospho log FC on the y-axis and Protein log FC on the x-axis, where the diagonal line (y = x) represented proportional changes between phosphorylation and total protein expression. For each phosphopeptide, deviation from this baseline was standardized as a z-score across all phosphopeptides, and statistical significance was determined by a two-tailed standard normal test (|z| > 1.96, p < 0.05). Significant phosphopeptides (|z| > 1.96) were further categorized by the magnitude of global protein change. Those with minor protein-level variation (|Protein log FC| < 0.25) were colored red (Phospho log FC > 0) or blue (Phospho log FC < 0), indicating phosphorylation-specific regulation independent of protein abundance. Conversely, phosphopeptides with substantial protein changes (|Protein log FC| ≥ 0.25) were colored orange, suggesting phosphorylation adjustments that may act compensatorily in response to altered protein expression levels.

### Ex vivo electrophysiology

Ex vivo whole-cell patch clamp recordings were performed to measure miniature excitatory postsynaptic currents (mEPSCs) from CA1 pyramidal neurons in male mice at postnatal weeks 6 to 9. Neurons were loaded with biocytin through the recording pipette for post hoc morphological analysis.

### Brain slice preparation

Mice were deeply anesthetized with isoflurane, and the brains were rapidly removed. Coronal sections (300 μm thick) were cut on a vibratome (VT1200S, Leica) in ice-cold (0 °C) sucrose-based dissection buffer containing (in mM) 75 sucrose, 76 NaCl, 25 NaHCO_3_, 2.5 KCl, 1.25 NaH_2_PO_4_(H_2_O), 25 D-(+)-glucose, 7 MgSO_4_(7H_2_O), and 0.5 CaCl_2_, continuously bubbled with 95% O_2_/5% CO_2_. Slices were first allowed to recover for 20 min in the same dissection buffer at 32 °C. They were then transferred to a holding chamber containing artificial cerebrospinal fluid (aCSF) at room temperature and kept there for at least 30 min before recording. The aCSF contained (in mM) 124 NaCl, 26.2 NaHCO_3_, 2.5 KCl, 1 NaH_2_PO_4_(H_2_O), 20 D-(+)-glucose, 1.3 MgCl_2_(6H_2_O), and 2.5 CaCl_2_, and was also bubbled with 95% O_2_/5% CO_2_.

### Whole-cell patch clamp recording

Slices were placed in a recording stage continuously perfused with aCSF at 26–28 °C. Recording glass pipettes were pulled from thin-walled borosilicate capillaries (30-0065, Harvard Apparatus) using a two-step vertical puller (PC-10, Narishige; first step 73.6 °C, second step 55–59 °C) and had a resistance of 2.9–4.6 MΩ. CA1 pyramidal neurons were visualized under differential interference contrast (DIC) optics on an upright microscope (B50WI, Olympus). CA1 pyramidal neurons were classified into deep and superficial sublayers based on soma position within the stratum pyramidale: neurons located closer to the stratum oriens were defined as deep, and those closer to the stratum radiatum as superficial based on visual inspection under DIC. Recordings and biocytin filling were performed in both sublayers in each slice. A gigaohm seal was formed between the pipette and the cell membrane, and the membrane was then ruptured to obtain the whole-cell configuration. Signals were acquired with a Multiclamp 700B amplifier and a Digidata 1440 digitizer (Molecular Devices), filtered at 2 kHz, and digitized at 10 kHz. Access resistance was monitored throughout each recording, and cells in which it exceeded 21 MΩ were excluded from analysis.

For mEPSC recording, pipettes were filled with an internal solution containing (in mM) 130 CsMeSO_4_, 10 TEA-Cl, 10 HEPES, 10 EGTA, 4 Mg-ATP, 0.3 Na-GTP, and 0.5 QX-314, supplemented with 0.25% (w/v) biocytin (hydrochloride; Sigma, B1758; pH 7.35; 295 mOsm before addition of biocytin). The aCSF contained 1 μM tetrodotoxin (TTX, Tocris) and 100 μM picrotoxin (PTX; Sigma) to block action potential-dependent synaptic currents and inhibitory postsynaptic currents, respectively.

mEPSCs were recorded for 2 min at a holding potential of −70 mV using Clampex 11.0.3 (Molecular Devices). Events were detected using the template search function in Clampfit 11.0.3 (Molecular Devices) with a template match threshold of 4 (in units of the noise standard deviation). A common template was applied to all recordings. Mean frequency and amplitude were calculated for each cell.

### Biocytin filling

After mEPSC recording, the amplifier was switched to current-clamp mode, and a continuous hyperpolarizing current of −50 pA (50 s per sweep, 12 sweeps) was injected twice to facilitate biocytin diffusion into the cell. The pipette was then withdrawn slowly to allow the membrane to reseal, and the slice was left in the recording stage for 2–3 min before fixation. Slices were then fixed in 4% PFA at 4 °C overnight.

### Biocytin labeling and dendritic morphology analysis

Following whole-cell recording, 300 µm-thick slices containing biocytin-filled neurons were fixed in 4% paraformaldehyde and stored at 4°C until staining. Slices were washed three times for 10 min in 1× PBS and blocked for 1 h in PBS containing 5% normal goat serum (NGS) and 0.2% Triton X-100. Slices were then incubated with an anti-NeuN antibody (MAB377, Millipore; 1:500) in blocking solution for two nights at 4°C. After five washes of 20 min each in PBS containing 5% NGS and 0.2% Triton X-100, slices were incubated overnight at 4°C with an Alexa Fluor 647-conjugated secondary antibody (115-606-071, Jackson ImmunoResearch; 1:500) together with Alexa Fluor 488-conjugated streptavidin (S11223, Invitrogen; 1:500). After final washes in the same buffer, slices were rinsed in PBS and mounted with mounting medium. Images were acquired on a confocal microscope (LSM900, Zeiss), and z-stack images spanning the entire dendritic arbor were acquired individually for each biocytin-filled neuron.

Biocytin-filled neurons were reconstructed in three dimensions using the Simple Neurite Tracer (SNT) plugin in ImageJ. Dendrites were traced manually, and apical and basal dendrites were analyzed separately. Total dendritic length was measured from the reconstructions, and Sholl analysis was performed on the same traced neurons using concentric spheres centered on the soma at 10 µm intervals to assess dendritic complexity as a function of distance from the soma. Neurons were assigned to the superficial or deep sublayer based on the position of their soma within the stratum pyramidale, and all analyses were performed by an experimenter blinded to genotype.

### Statistical analysis

Sample sizes were not predetermined by statistical methods or formal power analysis. Instead, sample sizes were determined based on our previous experience with this mouse model, sample sizes commonly used in similar neurobiological studies, animal availability, and ethical considerations to minimize animal use. In general, at least five biological replicate pairs were included per group, with most experiments using approximately eight pairs. Statistical analyses were performed using GraphPad Prism version 10.2.1 (GraphPad Software, San Diego, CA, USA). Outlier detection was performed using the ROUT (Robust Regression and Outlier Removal) method in GraphPad Prism. Data points identified as outliers were excluded from subsequent statistical analyses. Comparisons between two independent groups were performed using unpaired two-tailed Welch’s t-tests (with Welch’s correction). Comparisons among three or more groups with a single independent variable were performed using one-way analysis of variance (ANOVA) followed by Tukey’s or Šídák’s multiple comparisons test, as appropriate. When two independent variables were analyzed (e.g., genotype and time), two-way ANOVA followed by Tukey’s or Šídák’s multiple comparisons test was used, as appropriate. These tests were selected according to the number of groups being compared and the experimental design. Normality of each dataset was assessed using the Shapiro–Wilk test in GraphPad Prism. Results of all normality tests, including exact *P* values, are provided in Supplementary Table S16. A *P* value < 0.05 was considered statistically significant. Full statistical reports for all analyses are provided in Supplementary Table S17.

## Author Contributions

**U Suk Kim:** methodology, investigation, validation, formal analysis, visualization. **Yukyung Jun:** methodology, software, data curation, formal analysis, funding acquisition, visualization, writing - original draft. **Hyewon Kim:** methodology, software, data curation, formal analysis, visualization. **Yeji Yang:** methodology, data curation, investigation, formal analysis. **Muwon Kang:** methodology, investigation, formal analysis, visualization. **Ruiying Ma:** investigation, formal analysis. **Hyae Rim Kang:** investigation, formal analysis. **Sungjin Ju:** investigation, formal analysis. **Hoibin Jeong:** investigation, formal analysis. **Su Yeon Kim:** investigation, formal analysis. **Yoonhee Kim:** investigation, formal analysis. **Jun Young Oh:** investigation, formal analysis. **Eunjoon Kim:** funding acquisition, resources, writing - review & editing. **Kyoungmi Kim:** methodology, funding acquisition, resources. **Hyojin Kang:** software, data curation, investigation, funding acquisition, visualization, writing - original draft. **Jin Young Kim:** methodology, data curation, investigation, funding acquisition, resources, writing - original draft. **Kihoon Han:** conceptualization, supervision, funding acquisition, visualization, resources, writing - original draft, writing - review & editing.

## Acknowledgements

We thank the Korea Basic Science Institute (KBSI, Metropolitan Seoul Center) for providing access to 3D holotomography imaging (CC008).

## Ethics Statement

All animal maintenance and experimental procedures were approved by the Institutional Animal Care and Use Committee (IACUC) of Korea University College of Medicine (protocol no. KOREA-2024-0073). All procedures were conducted in accordance with the Laboratory Animal Act and the Animal Protection Act of the Republic of Korea, as well as the United States Public Health Service Policy on Humane Care and Use of Laboratory Animals.

## Data Availability Statement

The bulk RNA sequencing data of HEK293T cells and the single-nucleus RNA sequencing data of the mouse hippocampus have been deposited in the Gene Expression Omnibus (GEO) under accession numbers GSE317553 and GSE317598, respectively. The mass spectrometry proteomic and phosphoproteomic data have been deposited to the ProteomeXchange Consortium via the MassIVE partner repository with the dataset identifier PXD073375. All other data supporting the findings of this study are available within the article and its Supplementary Material, or from the corresponding authors upon reasonable request.

## Abbreviations

ACN: anhydrous acetonitrile
aCSF: artificial cerebrospinal fluid
Ago: Argonaute
ASOs: antisense oligonucleotides
ANOVA: analysis of variance
Brn2: Brain-specific homeobox/POU domain protein
2 Bsn: bassoon presynaptic cytomatrix protein
CA: Cornu Ammonis
CalB: Calbindin
CaMKIIα: calcium/calmodulin-dependent protein kinase II alpha
Cas9: CRISPR-associated protein 9
CRISPR: Clustered Regularly Interspaced Short Palindromic Repeats
Ctip2: COUP-TF-interacting protein 2
CUX1: Cut-like homeobox 1
CYFIP2: Cytoplasmic FMR1-interacting protein 2
Dab1: Disabled-1
DAPI: 4′,6-diamidino-2-phenylindole
Dcx: doublecortin
DEE65: developmental and epileptic encephalopathies 65
DEG: differentially expressed gene
DEP: differentially expressed protein
DG: dentage gyrus
Dgkβ: diacylglycerol kinase beta
DIC: differential interference contrast
DMEM: Dulbecco’s Modified Eagle Medium
EphA3: Eph receptor A3
ERK: extracellular signal-regulated kinase
FACS: fluorescence-activated cell sorting
FBS: fetal bovine serum
FC: fold change
FDR: false discovery rate
FMRP: fragile X messenger ribonucleoprotein
Gapdh: Glyceraldehyde-3-phosphate dehydrogenase
GEO: Gene Expression Omnibus
GFP: green fluorescent protein
GO: Gene Ontology
HCD: higher-energy collisional dissociation
HDR: homology-directed repair
HEK293T: human embryonic kidney 293T
IDT: Integrated DNA Technologies
KEGG: Kyoto Encyclopedia of Genes and Genomes
KO: knock-out
Lis1: Lissencephaly 1
Map2: microtubule-associated protein 2
Mark4: microtubule affinity-regulating kinase 4
mEPSCs: miniature excitatory postsynaptic currents
mTORC1: mechanistic target of rapamycin complex 1
Ndel1: nuclear distribution element-like 1
NeuN: Neuronal Nuclei
NGS: normal goat serum
OPC: oligodendrocyte precursor cell
Pafah1b1: platelet-activating factor acetylhydrolase 1b regulatory subunit 1
PAM: protospacer adjacent motif
PBS: phosphate-buffered saline
PCBP3: poly(rC)-binding protein 3
Pde4A: phosphodiesterase 4A
Prkcε: protein kinase C epsilon
Prox1: prospero homeobox 1
PTX: picrotoxin
PUM: Pumilio
PWK: postnatal week
RBFOX3: RNA binding fox-1 homolog 3
RBP: RNA-binding protein
ROUT: Robust Regression and Outlier Removal
Rplp2: ribosomal protein lateral stalk subunit P2
RRID: Research Resource Identifier (see scicrunch.org)
SDS: sodium dodecyl sulphate
sgRNA: single-guide RNA
snRNA-seq: single-nucleus RNA sequencing
SNT: Simple Neurite Tracer
ssODN: single-stranded oligo deoxynucleotide
Syn1: synapsin I
SynGAP1: synaptic Ras GTPase-activating protein 1
TBST: Tris-buffered saline with Tween 20
TEAB: triethylamonium bicarbonate
TFA: trifluoroacetic acid
TMT: plex tandem mass tag
TTX: tetrodotoxin
U2AF2: U2 small nuclear RNA auxiliary factor 2
UMAP: Uniform manifold approximation and projection
vGluT1: vesicular glutamate transporter 1
WAVE: Wiskott–Aldrich syndrome protein family verprolin-homologous protein
Wfs1: Wolfram syndrome 1
WRC: WAVE regulatory complex
WT: wild-type

