## Supplementary Figures 1-4 for "Imbalance between RNA-binding proteins and the WAVE regulatory complex underlies the toxic gain-of-function effects of the *CYFIP2* Arg87Cys variant in neurodevelopmental disorders"

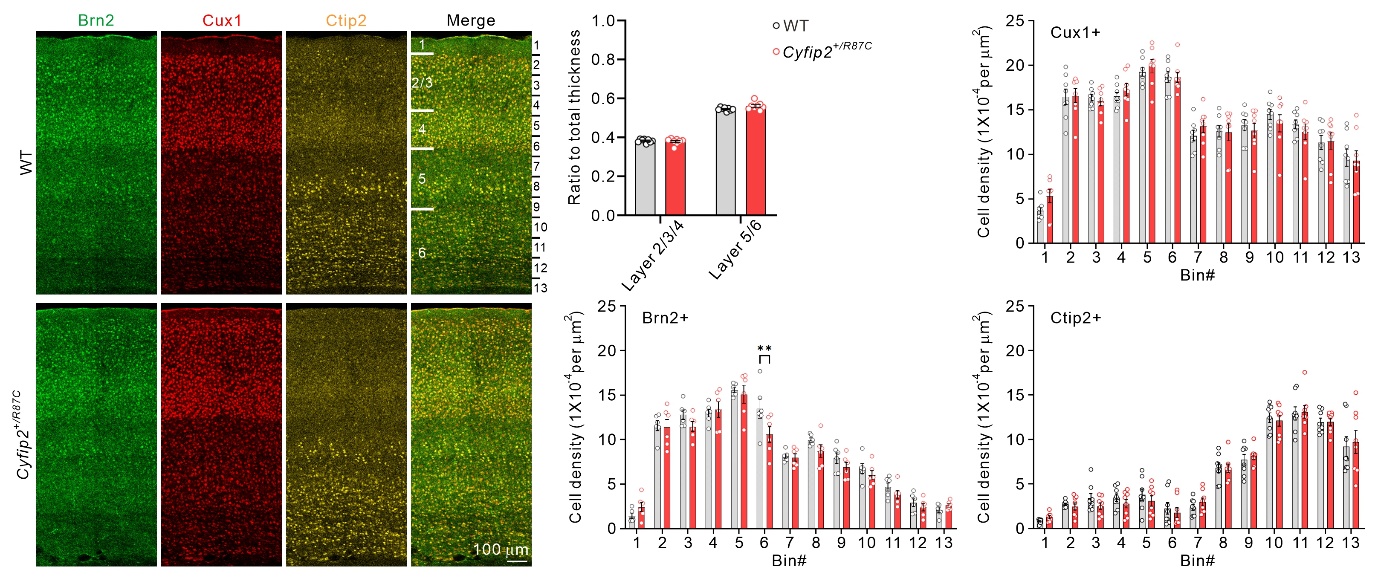

**Figure S1.** Normal neuronal layer organization in the somatosensory cortex of *Cyfip2^+/R87C^* mice. Representative fluorescent immunohistochemistry images stained with anti-Brain-specific homeobox/POU domain protein 2 (Brn2), anti-Cut-like homeobox 1 (Cux1), and anti-COUP-TF-interacting protein 2 (Ctip2) antibodies, together with corresponding quantification, reveal normal neuronal layer organization in the somatosensory cortex of *Cyfip2^+/R87C^* mice at postnatal week 7 (PWK 7) (n = 8 animals per genotype; two-way ANOVA with Šidák’s multiple-comparison test). Cell density was quantified in equally sized bins spanning the cortex from the pial surface to the white matter. A small but significant decrease in Brn2-positive cell density is observed only in bin #6 of *Cyfip2^+/R87C^* mice compared with wild-type (WT) mice. Data are presented as mean ± SEM. ***P* < 0.01.

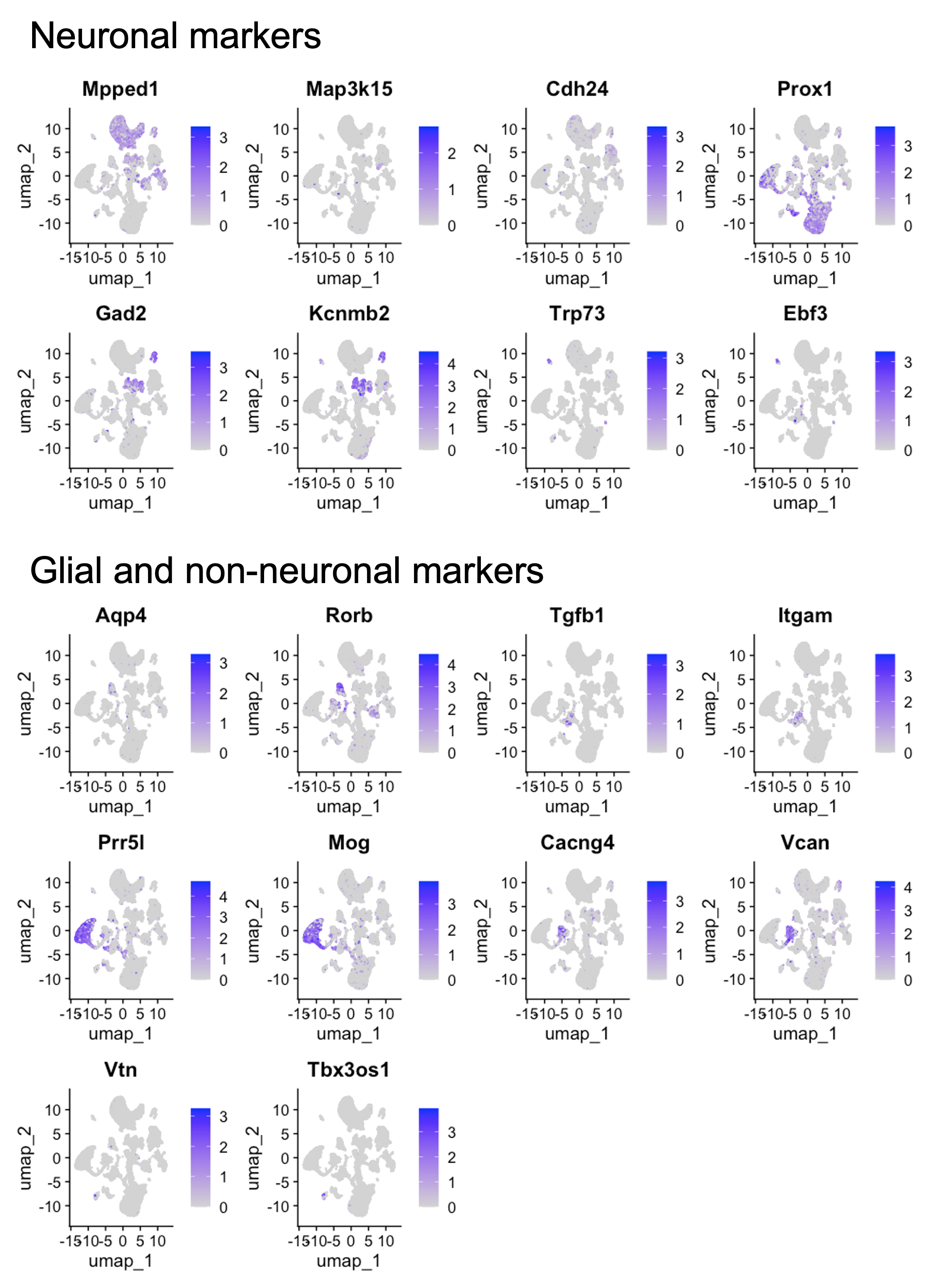

**Figure S2.** Representative marker gene expression used for cell type annotation in the single-nucleus RNA sequencing dataset. Feature plots show the expression patterns of representative marker genes across major neuronal populations and glial and non-neuronal populations. Neuronal markers include excitatory neuronal subtype markers corresponding to hippocampal subregions, namely Excitatory-CA1 (Mpped1), Excitatory-CA2 (Map3k15), Excitatory-CA3 (Cdh24), and Excitatory-DG (Prox1), as well as interneuron markers (Gad2, Kcnmb2) and Cajal–Retzius cell markers (Trp73, Ebf3). Glial and non-neuronal markers include astrocyte markers (Aqp4, Rorb), microglial markers (Tgfb1, Itgam), oligodendrocyte markers (Prr5l, Mog), oligodendrocyte precursor cell markers (Cacng4, Vcan), and fibroblast markers (Vtn, Tbx3os1).

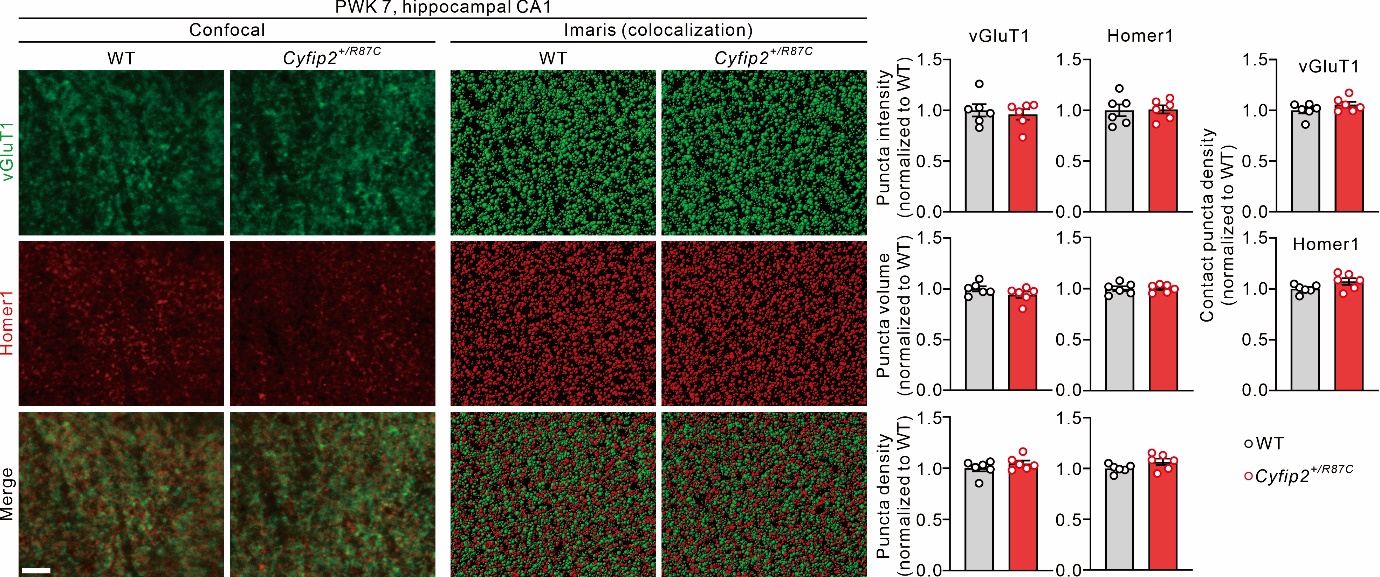

**Figure S3.** Normal excitatory synapse density in the hippocampal CA1 region of *Cyfip2^+/R87C^* mice. Representative fluorescent immunohistochemistry images stained with anti-vesicular glutamate transporter 1 (vGluT1) and anti-Homer1 antibodies and corresponding quantification reveal normal excitatory synapses in the dendritic layers of the hippocampal CA1 region of *Cyfip2^+/R87C^* mice at PWK 7 (n = 6 animals per genotype; unpaired two-tailed Welch’s t-tests). The intensity, volume, and density of vGluT1 and Homer1 puncta, and their colocalization, do not differ significantly between genotypes. Scale bar, 4 μm. Data are presented as mean ± SEM.

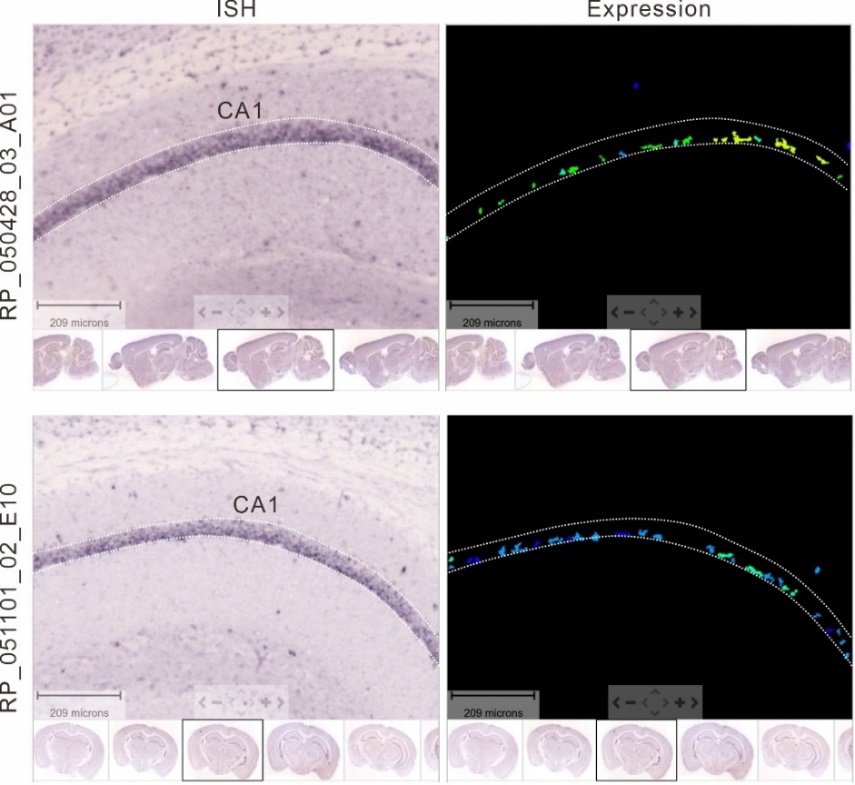

**Figure S4.** Preferential expression of *EphA3* mRNA in superficial-layer CA1 pyramidal neurons. *In situ* hybridization data from the Allen Brain Atlas show that *EphA3* mRNA is expressed in neurons in the superficial layer of the hippocampal CA1 region in the mouse brain. Data from two independent probes (A01 and E10) are shown. Expression images are displayed as pseudocolored maps of expression levels. Images were obtained from the Allen Mouse Brain Atlas (https://mouse.brain-map.org).
